# Metabolic modeling reveals the aging-associated decline of host-microbiome metabolic interactions in mice

**DOI:** 10.1101/2024.03.28.587009

**Authors:** Lena Best, Thomas Dost, Daniela Esser, Stefano Flor, Madlen Haase, A. Samer Kadibalban, Georgios Marinos, Alesia Walker, Johannes Zimmermann, Rowena Simon, Silvio Schmidt, Jan Taubenheim, Sven Künzel, Robert Häsler, Marco Groth, Silvio Waschina, Otto W. Witte, Philippe Schmitt-Kopplin, John F. Baines, Christiane Frahm, Christoph Kaleta

**Author notes:** shared second author, ordered alphabetically (Thomas Dost, Daniela Esser, Stefano Flor, Madlen Haase, A. Samer Kadibalban, Georgios Marinos, Alesia Walker, Johannes Zimmermann).

## Abstract

Aging is the predominant cause of morbidity and mortality in industrialized countries. The specific molecular mechanisms that drive aging are poorly understood, especially the contribution of the microbiota in these processes. Here, we combined multi-omics with metabolic modeling in mice to comprehensively characterize host–microbiome interactions and how they are affected by aging. Our findings reveal a complex dependency of host metabolism on microbial functions, including previously known as well as novel interactions. We observed a pronounced reduction in metabolic activity within the aging microbiome, which we attribute to reduced beneficial interactions in the microbial community and a reduction in its metabolic output. These microbial changes coincided with a corresponding downregulation of key host pathways predicted by our model to be dependent on the microbiome that are crucial for maintaining intestinal barrier function, cellular replication, and homeostasis. Our results elucidate microbiome–host interactions that potentially influence host aging processes, focusing on microbial nucleotide metabolism as a pivotal factor in aging dynamics. These pathways could serve as future targets for the development of microbiome-based therapies against aging.

Graphical abstract

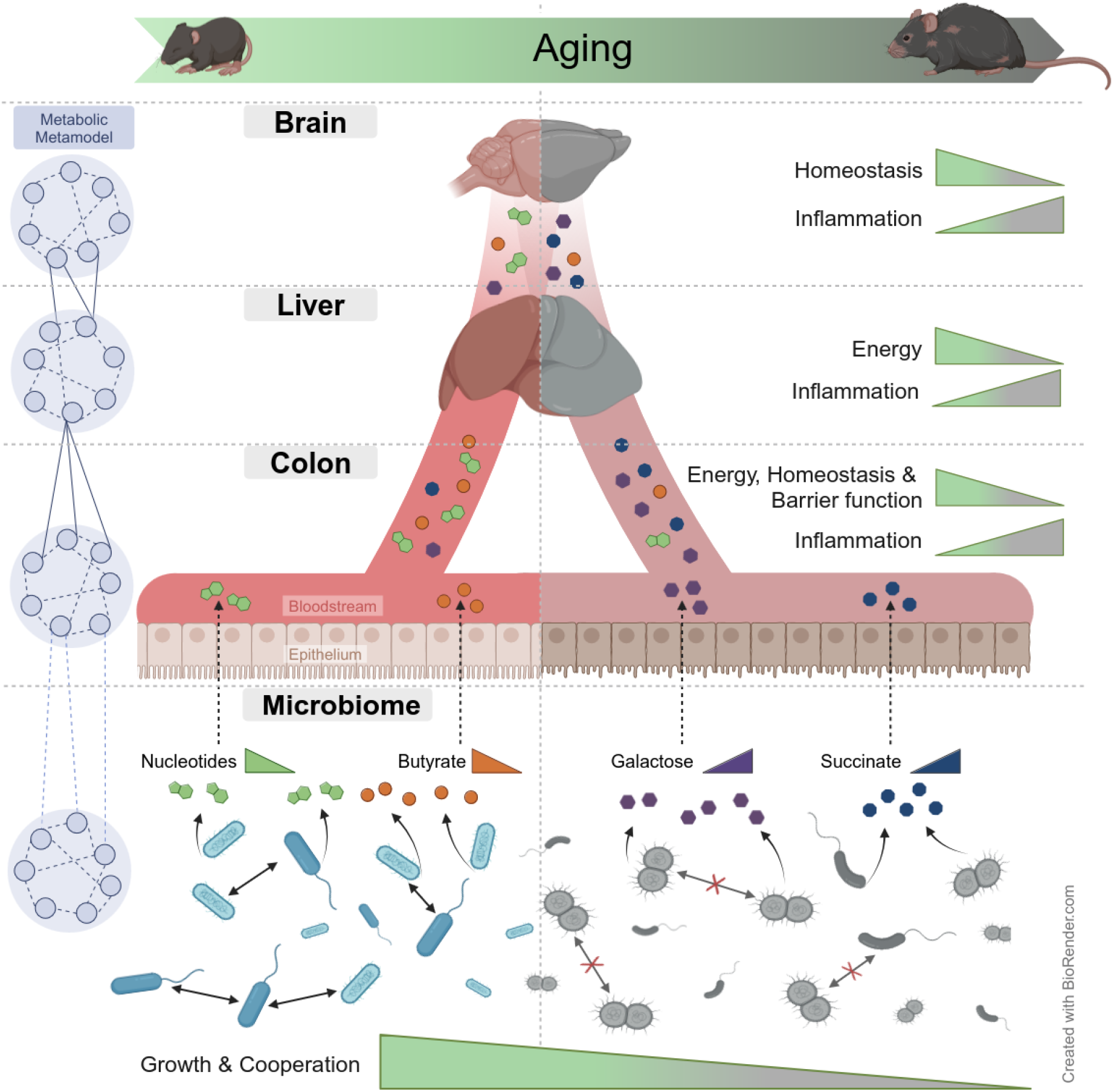

## Introduction

Aging and aging-related diseases are central contributors to morbidity and mortality in Western societies ^1^. Although research has identified specific hallmarks of aging ^2^ and revealed the conservation of aging-associated changes across species and tissues ^3^, the primary causative factors of aging remain elusive ^2,4^. The microbiome, comprising a diverse bacterial community that resides within and on host organisms, is gaining recognition for its significant interplay with host aging processes. It is implicated in many aging-associated physiological processes ^5–7^, showing notable shifts in its composition as the host ages and exhibiting strong correlations with aging-related phenotypes ^8^. Microbiome transfer experiments revealed that introducing old microbiota to young hosts induces aging-associated inflammation ^9^, whereas transferring young microbiota to old hosts extends lifespan ^10,11^ and reverses specific aspects of aging in animal models ^12^. Pathological changes in the host’s gastroenteric system, such as obstipation, constipation, and barrier dysfunction, are comorbidities of many aging-related diseases and often precede the manifestation of these diseases by many years ^13^. Moreover, the aging-associated loss of intestinal barrier function, which facilitates the translocation of living bacteria and their products into the bloodstream, is implicated as a driver of systemic inflammaging, a hallmark of aging characterized by a constant low-grade inflammation even without presence of a detectable pathogen ^9,14,15^.

However, it remains unclear which microbiome changes are causes of aging in the host and which are consequences ^5^. The primary reasons for this uncertainty are the high plasticity and complexity of the microbiota, which comprise dozens to hundreds of species ^16^, the low species-level conservation of microbes across human cohorts ^17^, and the myriad of metabolites through which the microbiota and host can interact ^18–20^. One approach to overcome this complexity is constraint-based metabolic modeling ^21,22^; this method builds on *in silico* representations of the metabolic networks of individual species—so-called genome-scale metabolic networks and allows the prediction of metabolic fluxes in individual species ^21^ or entire communities ^22,23^. This approach enables the integration of different types of omics datasets to derive context-specific metabolic networks (i.e., networks representing the metabolic state of particular tissues or cells) ^24^. Therefore, several studies have used constraint-based metabolic modeling to investigate changes in microbiome–host interactions in various diseases ^25,26^ and identify specific microbial processes linked to therapeutic response ^23,27^.

In this study, we used tissue transcriptomic, metagenomic, and metabolomic data to elucidate the metabolic mechanisms through which the gut microbiota could contribute to host aging. We extensively characterized microbiome–host interactions at the level of global associations between host transcript levels and microbiome functions and then focused on metabolic interactions using an integrated metabolic model of the host and the microbiota. Our results revealed many known interactions between the host and the microbiota and postulated numerous novel ones. Next we investigated how these interactions change in the context of aging. We observed a considerable reduction in microbiome metabolic activity with age, which seemed to be driven by substantial changes in within-microbiota ecological interactions. We then linked aging-associated changes between the host and the microbiota and found that aging-regulated genes were highly enriched for microbiome-dependent genes and model-predicted microbiota-dependent host functions, showing a pronounced suppression with age. These findings indicate that the microbiome is a major contributor to aging-associated metabolic decline, which we also observe at the metabolome level and thereby pinpoints metabolic pathways through which the microbiome may influence aging in the host.

## Results

### Taxonomic and functional description of the mouse microbiome

We studied the effects of aging in 52 male wild-type C57BL/6J/Ukj mice, separated into five age groups between 2 to 30 months old; 2 months corresponds to the stage after the transition from adolescence to adulthood, and 30 months marks late aging, with a 5% survival rate ^28^. We obtained transcriptome sequencing data for the colon, liver, and brain of these mice, as well as shotgun (167 Gbp) and long-read sequencing data (13.7 Gbp) for fecal samples, which we used to reconstruct 181 metagenome-assembled genomes (MAGs) of bacteria comprising their gut microbiome (Fig. 1A, see Supplementary Figs. 1A–B and Methods). The combined genome size over all 181 MAGs summed up to 367 Mbp and was distributed across 16,656 scaffolds. Taxonomic classification with the genome taxonomy database toolkit (GTDB-Tk) ^29^ assigned 175 MAGs to known taxa by closest phylogenetic placement or average nucleotide identity value (with the prefix “GCA_” or “GCF_” in Fig. 1A), whereas 6 MAGs could not be assigned (prefixed “NEW_” in Fig. 1A). Of those 181 MAGs, 25 were considered high-quality drafts according to established criteria ^30^, and the rest were considered medium-quality drafts. Notably, we used more stringent cutoffs (≥80% completeness and ≤10% contamination) than those suggested by Bowers et al. ^30^ for medium-quality MAGs to require less gap-filling and thus obtain more reliable metabolic models for downstream analysis (Fig. 1A). Most of the MAGs were attributed to the phyla Bacillota (previously Firmicutes; *n* = 97) and Bacteroidota (*n* = 65). The reconstructed genomes from rarer phyla included Pseudomonadota (previously Proteobacteria; *n* = 9), Cyanobacteriota (previously Cyanobacteria; *n* = 4), Campylobacterota (*n* = 3), Deferribacterota (*n* = 1), Desulfobacterota (*n* = 1), and Verrucomicrobiota (*n* = 1). Regarding overall abundance, the most abundant MAGs, with a coverage depth >1%, belonged to Bacteroidota in the families Bacteroidaceae (*n* = 5) and Muribaculaceae (*n* = 12). Only three MAGs in the Lachnospiraceae family (Bacillota) and one in Akkermansiaceae (Verrumicrobiota) surpassed a 1% coverage depth (Fig. 1A; Supplementary Table S1.2, Supplementary Fig. 1F). The genome sizes of the MAGs ranged from 0.9 to 6.7 Mbp (Fig. 1A).

To functionally annotate the assembled MAGs, we used gapseq ^31^ to reconstruct their corresponding genome-scale metabolic networks. We explored their metabolic diversity with principal component analysis (Fig. 1B). Principal component (PC) 1 mainly separated the metabolic models by the completeness score (*R*^2^ = 0.15) of the underlying MAGs, but it also separated them by the taxonomic rank “order” (*R*^2^ = 0.87). The completeness of the MAGs significantly impacted the prevalence of pathway gaps within the reconstructed models. Consequently, the occurrence of such gaps (*R*^2^ = 0.55) and the sizes of the models (*R*^2^ = 0.84) or genomes (*R*^2^ = 0.58) partially accounted for the observed differentiation along the first two PCs. PC2 separated the metabolic models by the phylum, GC content (*R*^2^ = 0.29), and contamination score (*R*^2^ = 0.06) of the underlying MAGs. The contamination score was inversely associated with the number of successfully recovered tRNA genes in the MAGs (*R*^2^ = 0.17).

**Figure 1:**
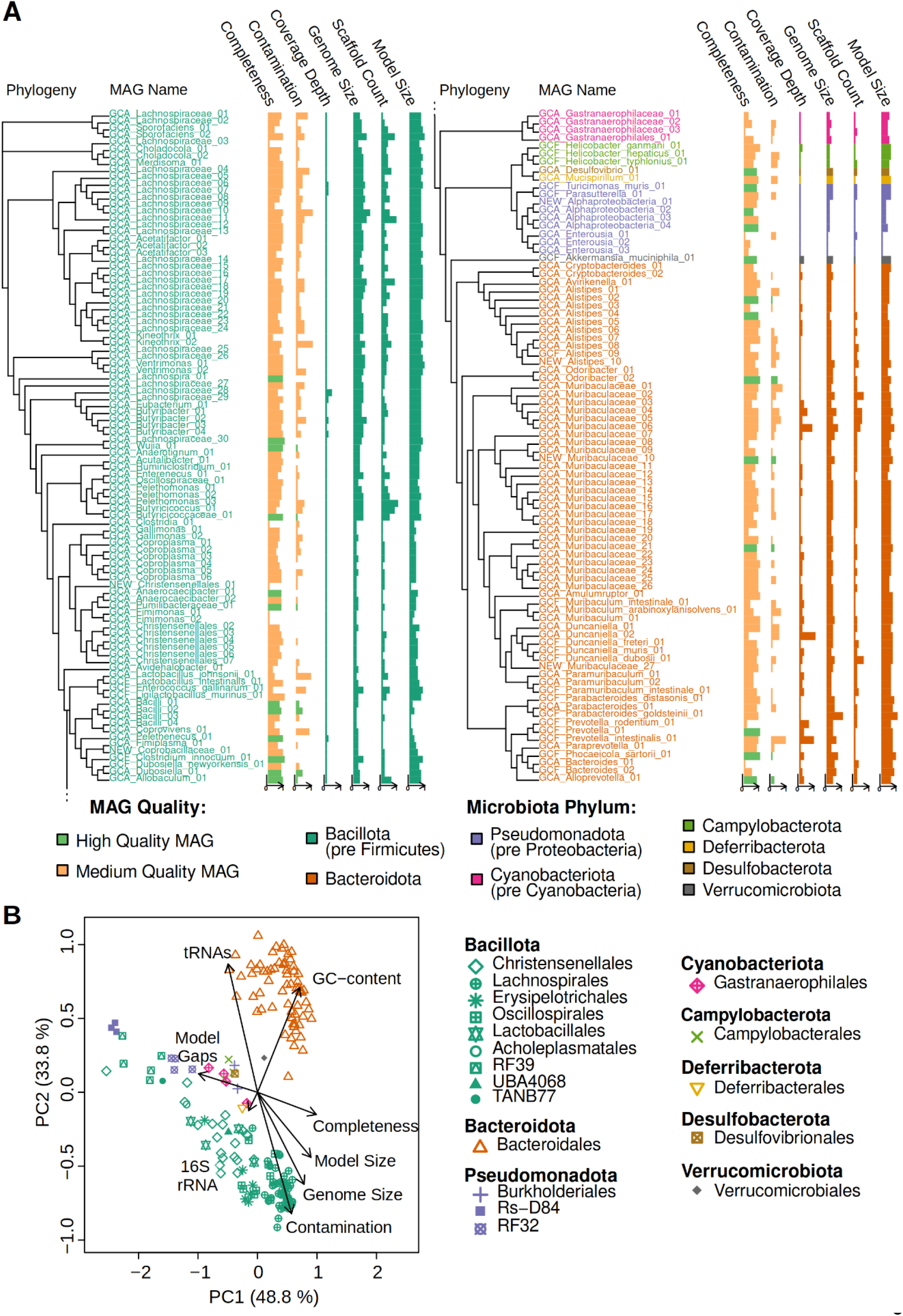
Mouse microbiome and metabolic model characterization. **A** Phylogenetic tree of the 181 MAGs (split after 97 Bacillota MAGs for clarity), with the MAGs assigned names, completeness (81.5%–100%), and contamination (0%–9.8%), as well as the total coverage depth across the entire cohort (26.9–2846.5), the draft genome sizes (0.9–6.7 Mbp), their MAG fragmentation (5–554 scaffolds), and the number of reactions in the reconstructed metabolic models (726–1985). See Supplementary Table S1.2 for further metadata information. **B** Principal component analysis of the metabolic models of the mouse microbiota. Inter-model distances are based on the Horn–Morisita dissimilarity index of the functional content of each model (see Methods). Metadata associations to PCs are overlaid as arrows. The shapes denote the taxonomic rank “order,” whereas the coloring of the symbols in **B** follows the coloring for the phyla in **A**.

### Microbiome functions correlate widely with host transcripts across tissues

After reconstructing the metabolic models of the bacterial species of the mice, we first conducted a global assessment of which host functions were associated with microbial metabolic activity independent of their association with age. This was achieved by correlating the occurrence of metabolic reactions within the microbiota with host gene expression, adjusting for age with partial Spearman’s rank correlation. Considering associations with a strong correlation coefficient and a false discovery rate (FDR)-adjusted *p*-value of ≤ 0.1, we identified 12,732 strongly correlated microbiome reactions and host genes for the colon, along with 3,425 for the liver and 2,499 for the brain. Enriching these correlated features with gene ontology (GO) ^32^ biological processes (host genes), and metabolic subsystems (microbiome reactions), respectively, we obtained 1,377 pairs of host–microbiome-associated processes for the colon, 283 for the liver, and 167 for the brain; we further summarized these with level 2 GO biological processes and MetaCyc ^33^ superpathways (Fig. 2A–F; Supplementary Tables S2.1–S2.4).

**Figure 2:**
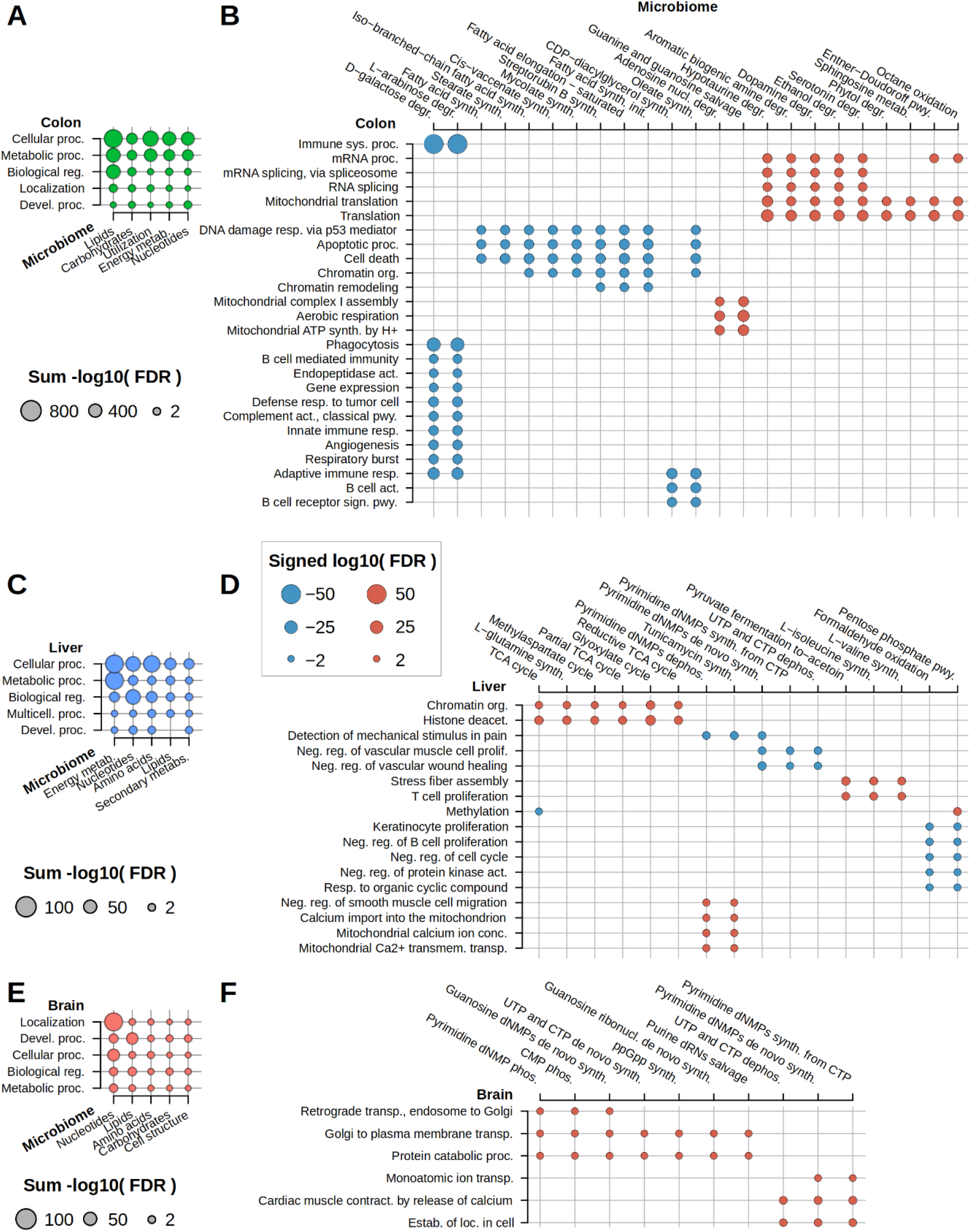
Correlation-derived host–microbiome interactions. **A, C, E** Summaries of interaction scores (sum of −log_10_ FDRs) between host level 2 GO biological processes and microbiome super-pathways for the colon (**A**), liver (**C**), and brain (**E**). **B, D, F** Interactions between host biological processes and microbiome metabolic subsystems for the colon (**B**), liver (**D**), and brain (**F**). Dot size represents the log_10_ FDR-corrected p-values from over-representation tests, and the color represents either negative (blue) or positive (red) associations. The processes with the most significant associations (colon FDR ≤1·10^−10^, liver FDR ≤1·10^−4^, and brain FDR ≤1·10^−3^) with at least two interactions are shown (see Supplementary Tables S2.1–2.4 for the complete list).

The most strongly correlated host functions for the colon involved innate and adaptive immune processes and protein processing (Figs. 2A and 2B; Supplementary Table S2.1). These included a negative correlation between host immune system processes and microbial galactose and arabinose degradation pathways. Moreover, we observed strong positive correlations between microbial purine metabolism and mitochondrial respiration in the host. Furthermore, we found that microbial pathways involved in lipid metabolism were correlated with host processes involved in tissue homeostasis, such as DNA damage responses and cell death. For the liver, we detected strong correlations between the central metabolic pathways of the microbiota and chromatin organization in the host and between T-cell proliferation and microbial branched-chain amino acid metabolism (Figs. 2C and 2D; Supplementary Table S2.2). For the brain, we found strong correlations between protein catabolic processes and microbial nucleotide metabolism (Figs. 2E and 2F; Supplementary Table S2.3).

### Metabolic metaorganism models reveal widespread metabolic interactions between host and microbiota

After extensively characterizing the associations between host transcripts and microbiome metabolic functions, we next aimed to gain a more mechanistic understanding of the underlying pathways mediating those associations at the metabolic level with an integrated metabolic metamodel of the host and the microbiome. In this metamodel, the host is represented by three different tissues (colon, liver, and brain) connected through the bloodstream and interacting with the microbiome through the gut lumen (Fig. 3A). Each host tissue is represented by a unique instance of the human metabolic reconstruction Recon 2.2 ^34^, whereas the microbiome is represented by a combined model including all the metabolic reactions occurring in at least one bacterial metabolic model reconstructed from the MAGs (Fig. 3A). Starting from the generic metamodel, context-specific metabolic metamodels representing the metabolic state of each mouse were built by inputting tissue transcriptomic data and metagenomic abundance data into fastcore ^35^ (Supplementary Table S3.1).

**Figure 3:**
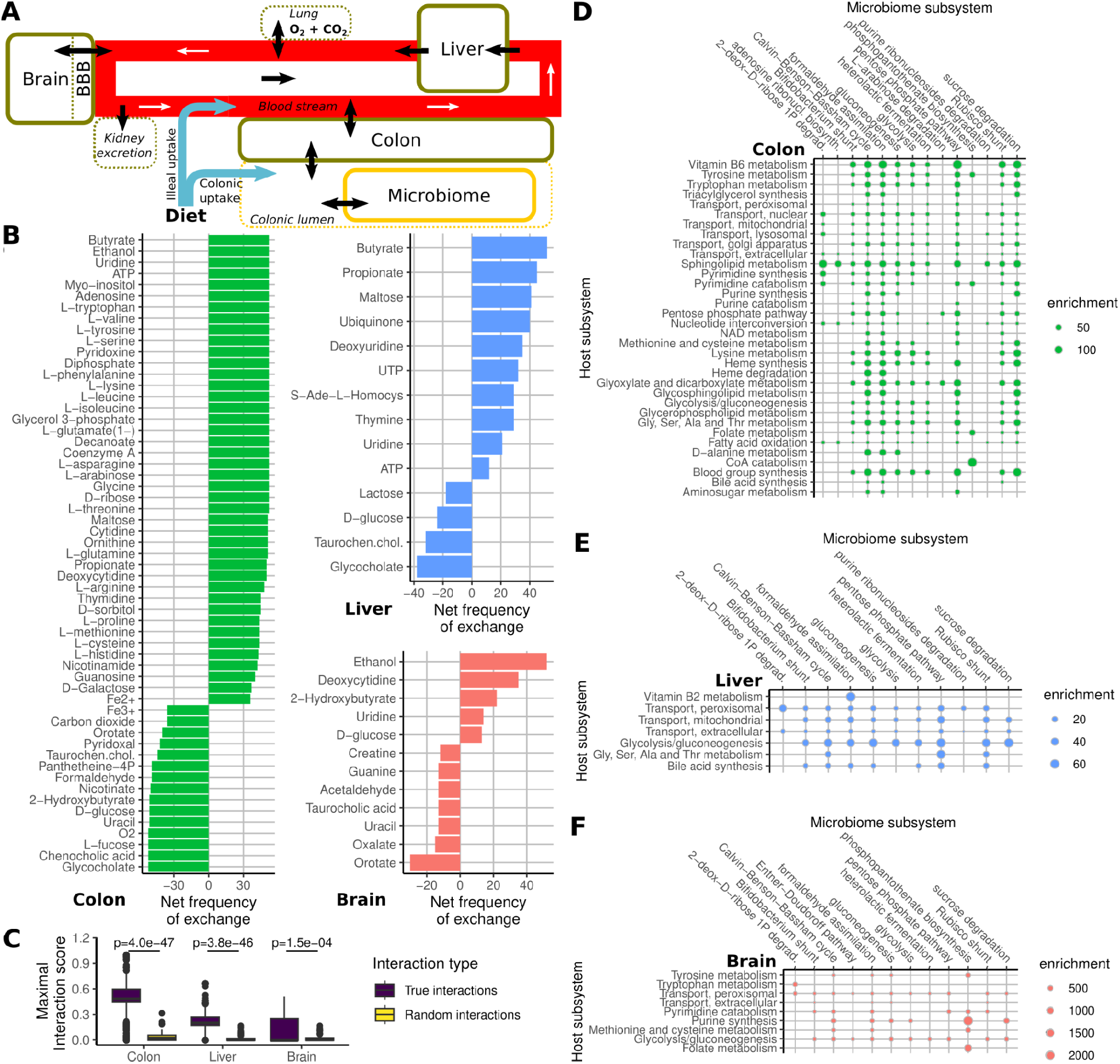
Model-predicted host–microbiome interactions. **A** Schematic structure of the metamodels. Solid borders indicate compartments of the metamodel with individual metabolic reactions and metabolites. Black arrows indicate metabolite exchanges between compartments and their surroundings (i.e., the gut lumen and bloodstream). Dashed borders indicate compartments only represented by inflow or outflow reactions. White arrows indicate the direction of metabolic exchanges along the bloodstream. BBB, blood-brain barrier. **B** Frequency of microbiome dependence on metabolite import (positive numbers) and export (negative numbers) across organs. Metabolites with the highest frequency of exchange across 52 models are shown (see Supplementary Table S3.2 for the complete list). **C** Maximal interaction scores for pairs of host–microbiome-associated processes versus 100 randomly selected pairs of host genes and microbiome reactions for each tissue (see Methods). The p-values indicate the significance of the differences according to the Wilcoxon rank-sum test. **D–E** Subsystem enrichment of model-predicted interactions between host and microbiome reactions. Only microbiome subsystems interacting with at least two host subsystems with an enrichment p-value <10^−4^ are shown (Fisher’s exact test; see Supplementary Tables S3.3–S3.5 for the complete list).

To explore the extent to which the metamodel could reconstitute known host–microbiome interactions at the metabolic level, we used it to predict mutual metabolic exchanges between the host and the microbiota (Fig. 3B; Supplementary Table S3.2). Among the metabolites identified in the colon, several involved known host-derived compounds, such as bile acids and fucose, a component of mucins ^36^. The microbiota provided the short-chain fatty acids propionate and butyrate as well as tryptophan and other essential amino acids, as previously documented ^37^. Moreover, we observed that the microbiota produced many nucleotides, including nucleotide derivatives such as NAD and coenzyme A. For the liver, we observed metabolic interactions involving the provision of primary bile acids to the microbiota and microbial production of nucleotides and short-chain fatty acids. For the brain, we observed that the host was provided with the microbial fermentation product ethanol and several pyrimidines. As observed in the colon, the host provided the pyrimidine precursor orotate and the nucleotide degradation product uracil to the microbiota, and the microbiota provided uridine and deoxycytidine in return.

To elucidate the underlying metabolic pathways between the host and the microbiota that mediate the extensive associations we identified between host transcripts and microbiome metabolic functions, we sampled elementary flux modes (EFMs) corresponding to minimal metabolic pathways ^38^ in the metamodel with EFMSampler ^39^. Each host reaction was defined as an indicator reaction through which EFMs were sampled. By recording the frequency at which microbial reactions occurred in the EFMs of a host indicator reaction, we obtained an interaction matrix of the frequency at which microbiome reactions occurred in the pathways sampled for individual host indicator reactions. This matrix indicates direct dependencies of host reactions on microbial reactions. Reassuringly, when comparing the predicted interaction scores of pairs of host–microbiome-associated processes (cf. Fig. 2B, D, F) with randomly sampled pairs, we found significantly higher interaction scores among the correlated pairs of host–microbiome-associated processes for the liver and colon. However, the interaction scores of the randomly sampled interaction pairs were slightly higher for the brain (Fig. 3C; see Methods).

To further functionally characterize the interaction matrix, we performed enrichments for host and microbial metabolic subsystems (Fig. 3D–F). We found that the most extensive interactions in the colon involved host pathways associated with energy metabolism, nucleotide metabolism, vitamin metabolism, and amino acid metabolism (Fig. 3D). Consistent with the correlational analyses, the colon depended on fermentation products, nucleotide metabolism, and vitamin biosynthesis pathways of the microbiota. In the liver, energy-producing pathways and bile acid synthesis were prominent on the host side, whereas the microbial side exhibited enrichment in microbial fermentation pathways (Fig. 3E). In the brain, microbiome-dependent host reactions exhibited enrichment in nucleotide metabolism, folate metabolism, and the metabolism of various amino acids, including the precursors of essential neurotransmitters, such as tryptophan and tyrosine (Fig. 3F). Although most host–microbe interactions were relatively generic, relying on basic microbial metabolic functions (such as glycolysis and fermentation), we also identified specific interactions, such as colonic nucleotide interconversion dependent on microbial ATP synthesis and colonic coenzyme A catabolism reliant on microbial production of phosphopantothenate, a coenzyme A precursor.

### Aging is associated with a profound loss of microbiome metabolic activity

After extensively characterizing host–microbiome interactions in the mouse cohort, we explored how these interactions change in the context of aging. First, we identified aging-associated changes in the microbiota. Consistent with previous reports in mice ^40,41^, we observed that increased age was associated with a decrease in the abundance of species in the phylum Bacillota and an increase in the abundance of most species in the phylum Bacteroidota (Fig. 4A). To obtain a better functional understanding of these species-level changes, we used community flux balance analysis (FBA) ^23^ to predict microbial metabolic activities on the basis of dietary and bacterial abundance information (see Methods). We determined the associations between metabolic activities and age and summarized them at the pathway level. We mainly observed negative associations (Fig. 4B) involving many biosynthetic pathways essential for bacterial replication, such as synthesizing amino acids, nucleotides, vitamins, and cell wall components. Similarly, for metabolic interactions between the microbiota and the host, we mainly observed strong reductions in both the consumption and production of metabolites (Fig. 4C), including microbial production of the anti-inflammatory short-chain fatty acid butyrate, and increased production of selected metabolites, such as pro-inflammatory succinate ^42^. Consistent with a generally reduced microbial metabolism, we also found that model-predicted and metagenomics-derived microbial growth rates decreased considerably with age (Fig. 4D) and were strongly correlated (Supplementary Fig. 4C).

**Figure 4:**
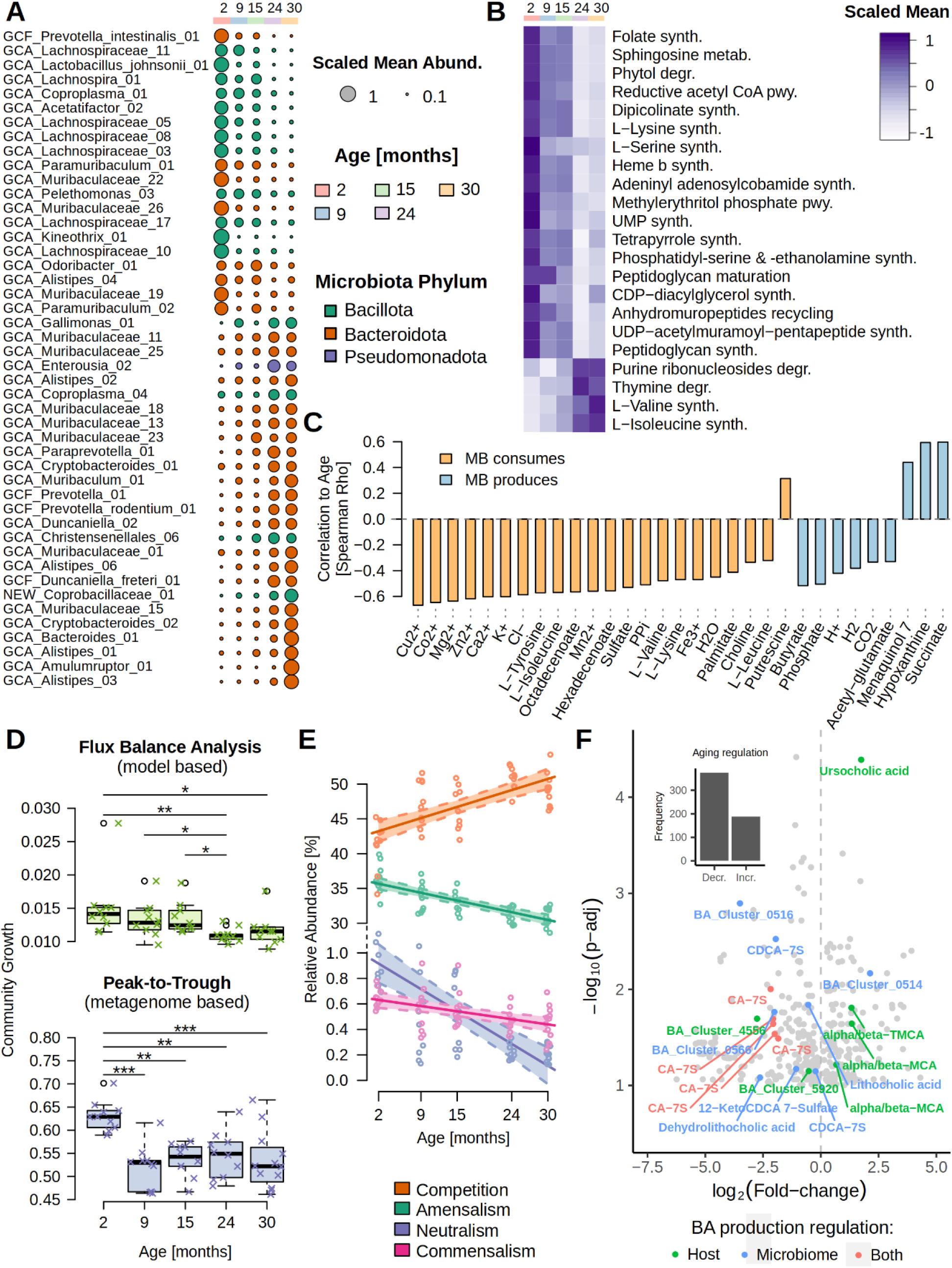
Microbiome alterations associated with host age. **A** Aging-associated changes in MAG abundance. **B** Aging-associated changes in microbiome internal reaction fluxes at the subsystem level. **C** Aging-associated changes in the host–microbiota metabolic exchange. **D** Comparison of community growth rates derived from FBA or peak-to-trough ratio. **E** Aging-associated changes in model-predicted ecological interactions in the microbiota. **F** Aging-associated changes in fecal metabolite concentrations in mice. All metabolites that were significantly associated with age (FDR-adjusted p ≤ 0.1) are shown. For clarity, only log_2_ fold changes in the range [−7.5, +5] and −log_10_ p-values of <4.5 are shown (no annotated metabolites were outside this range; see Supplementary Table S4.12 for the complete data). The origin of bile acids is indicated. “Both” refers to cholic acid-7-sulfate produced by the host but whose production is regulated by the microbiota ^48^. Metabolite names starting with “BA_Cluster” indicate bile acids that have not been fully resolved. Abbreviations: CA-7S, cholic acid-7-sulfate; CDCA-7S, chenodeoxycholic acid-7-sulfate; MCA, muricholic acid; TMCA, tauromuricholic acid. Asterisks indicate the significance of differences: *, p ≤ 0.05; **, p ≤ 0.01; ***, p ≤ 0.001.

To gain insight into the potential microbiome-intrinsic causes of the observed aging-associated reduction in microbiome metabolism, we used community FBA ^43^ to predict the frequencies of ecological interactions (see Methods). We observed a significant decrease in amensal, commensal, and neutral interactions at the expense of increased competitive interactions (Fig. 4E). These shifts in community interactions were also observed at the level of individual microbial ecological strategies derived from the universal adaptive strategies theory framework ^44,45^, indicating a shift in the community toward the dominance of ruderals, which are less diverse metabolically and interact less frequently with other species (Supplementary Fig. 4B). To further explore the predicted loss of microbiome metabolic activity with age, we performed an untargeted metabolomics analysis of fecal samples from an independent cohort of 82 mice across all age groups. We determined the correlation between the abundance of identified metabolic clusters and age and found that 374 out of 561 clusters (67%) showed significant downregulation (FDR-adjusted *p* ≤ 0.1), further supporting age-associated repression of microbial metabolism (Fig. 4F). Within this dataset, we specifically annotated bile acids using reference standards because of their previously documented role in host aging ^46^ and the possibility of distinguishing between host-regulated and microbiome-regulated bile acids. Consistent with the reduction in microbiome metabolic activity with age, we found that the concentrations of host-regulated bile acids were significantly increased (four out of six clusters). By contrast, the concentrations of microbiome-regulated bile acids were mostly reduced (seven out of eight clusters). Intriguingly, clusters annotated as cholic acid-7-sulfate were exclusively downregulated with age. Although this bile acid is known to be produced by the host ^47^, a previous study demonstrated that its synthesis in the host is regulated by microbial production of lithocholic acid and is strongly downregulated in antibiotic-treated animals ^48^. Further annotation of metabolic clusters identified that other microbiome-regulated metabolites, including valine/betaine, nicotinamide, enterolactone, and 3-hydroxykynurenine, were downregulated with age (Supplementary Table S4.12). Moreover, we found an increase in the pro-inflammatory microbial metabolite D-galactose, for which we observed a strong association with host immune processes in the colon (Fig. 2B), although this aging-associated increase was only significant before FDR correction (Supplementary Fig. 4D).

### Conserved aging-associated changes across mouse tissues

We conducted a differential gene expression analysis followed by GO enrichment of the significantly changed transcripts to gain insights into aging-associated changes in the host. Consistent with our previous observation of a signature of aging conserved across species and tissues ^3^, we observed a strong conserved signature of aging across the colon, liver, and brain in our mice. Overall, we observed increased activity for inflammatory and immune processes in aged mice across all three tissues (inflammaging) and decreased proliferative potential in the colon and brain. Other examples of aging-induced processes shared across all tissues were lipopolysaccharide (LPS), defense, immune, and inflammatory responses, along with the formation of blood vessels (Fig. 5A). Nervous system development was downregulated in both the aged colon and brain, which appeared to occur at an earlier age in the colon (9–15 months) than in the brain (15–24 months). The downregulation of cell division was unique to the aged colon (Fig. 5B). In the liver, processes related to mitochondrial energy production, protein translation, and assembly showed age-related decreases, whereas immune processes, signaling pathways, and proliferation processes (such as cell migration, cell proliferation, and extracellular signal-regulated kinase cascades) showed age-related increases (Fig. 5C). In the brain, gene expression related to microglial cell activation and LPS-related and pattern recognition signaling pathways showed age-related increases, whereas many learning, memory, synaptic plasticity, and synaptic signaling pathways showed age-related decreases (Fig. 5A, D). Focusing on gene-level conservation of aging-associated regulation, we found 157 transcripts that were consistently downregulated with age and 526 transcripts that were consistently upregulated with age. The significantly induced genes were enriched for regulation of interleukin (IL)-1b, IL-2, IL-4, IL-6, IL-10, IL-12, IL-17, tumor necrosis factor (TNF), interferon (IFN)-α, IFN-β, IFN-ɣ, C-X-C motif chemokine ligand 2 (CXCL2), and immunoglobulin and cytokine production (Supplementary Fig. 5; Supplementary Tables S5.7–S5.9). Because we found an aging-associated increase in microbial production of the pro-inflammatory metabolite succinate ^42,49^ (Fig. 4C), we further explored aging-associated changes in succinate-related genes. The sodium succinate transmembrane antiporter solute carrier family 13 member 3 (Slc13a3) was significantly downregulated in the colon and the brain of aged mice, whereas it was upregulated in the liver. Conversely, sirtuin 5 (Sirt5), a regulator of mitochondrial energy production and succinate dehydrogenase ^50^, showed the opposite regulation patterns (upregulation in the colon and brain and downregulation in the liver; Supplementary Tables S5.1–S5.3).

**Figure 5:**
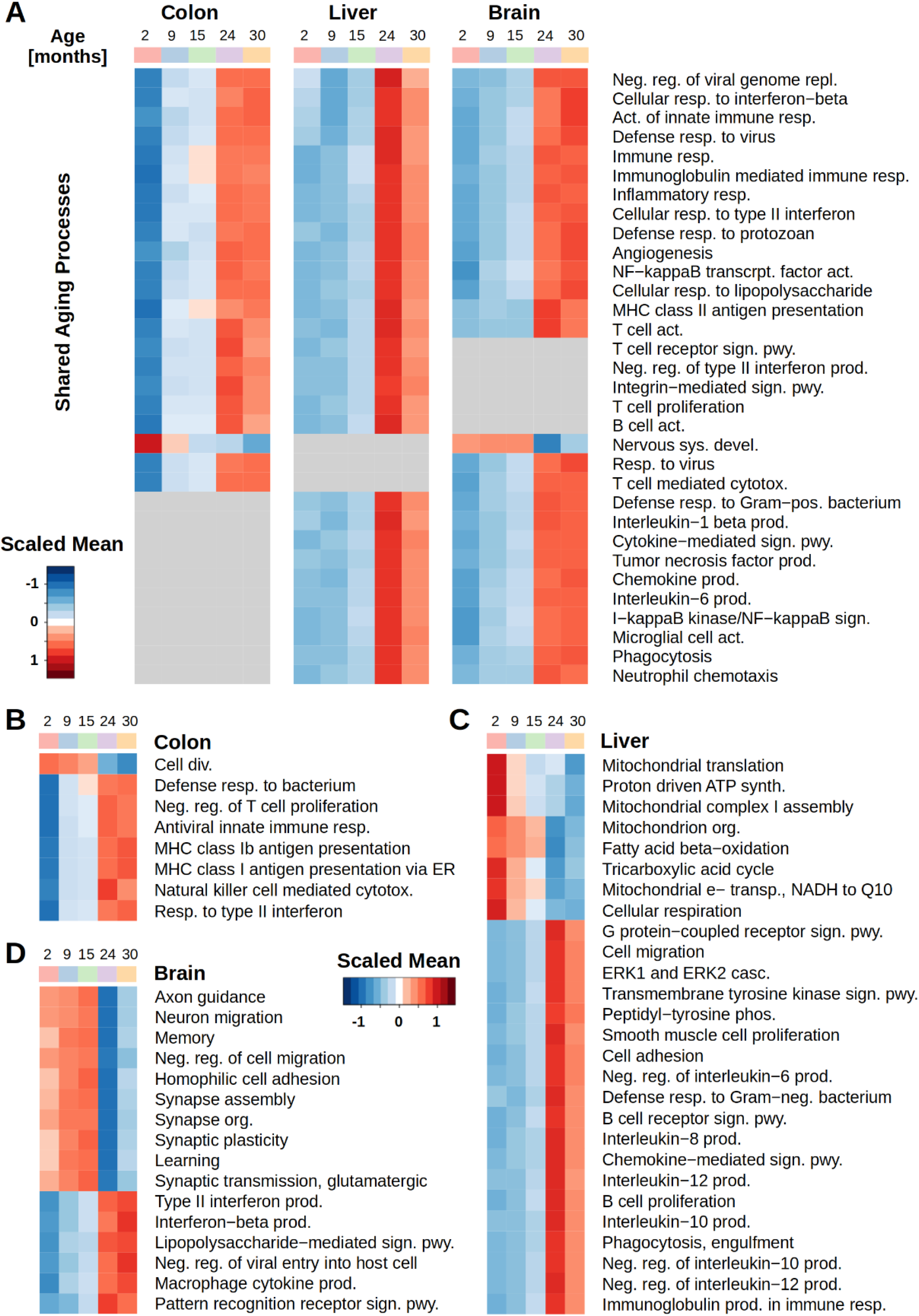
Aging-associated transcriptomic changes across host tissues. Differentially expressed genes were enriched for GO biological processes. The average expression abundance of all enriched features for each GO term is plotted stratified by age group and organ. **A** Enriched GO terms shared by at least two organs. **B–D** Enriched GO terms unique to each organ. The enrichment FDR-adjusted p-value cutoff for displaying a GO term was 10^−4^ for the colon and 10^−6^ for the liver and brain.

### Aging-associated decline in microbial metabolic activity impacts host functions

Next, we investigated how the aging-associated loss of microbiome metabolic function potentially impacted host functions. Among aging-regulated host genes, we found a highly significant enrichment of microbiome-correlated transcripts across all three tissues (Fig. 6A). This correlation was not driven by the indirect correlation of host and microbiome functions with age; we explicitly corrected for age when obtaining microbiome-correlated host genes (see Methods). Along with the loss of microbiome metabolic function with age, we also found a significant decrease in the frequency of significant correlations between host transcripts and microbiome metabolic functions with age (Supplementary Fig. 6A–E). GO biological process enrichment of genes associated with both aging and the microbiome revealed extensive aging-regulated and microbiome-associated processes in the colon but fewer in the liver and brain (Fig. 6B, Supplementary Fig. 6F, G; Supplementary Tables S6.5–S6.8). Notably, tissue homeostasis and organ regeneration processes were downregulated in the colon with age but positively correlated with microbial metabolic pathways. Conversely, aging-induced processes, primarily defense, inflammatory, and immune responses, were negatively associated with microbial metabolism (Fig. 6B). Brain development was negatively correlated with microbial metabolism and down-regulated with aging (Supplementary Fig. 6G). Further analysis of microbial subsystems identified a reduction in glycolysis, nucleotide synthesis, and D-galactose degradation with age, which were mostly inversely related to host gene expression (Fig. 6C). Given the observed reduction in interactions at the level of associations between host transcript levels and microbiome metabolic functions, we next aimed to identify the metabolic pathways linking the host and microbiota that underlie these associations. To achieve this, we defined aging-regulated metabolic modules within the metamodel’s metabolic reactions with the reaction-level interaction matrix between host and microbiome reactions obtained by EFM sampling. Metabolic modules were determined according to sampled EFMs (cf. Fig. 3D–F), selecting reactions present in at least 20% of the EFMs for each indicator reaction (see Methods). Thus, each metabolic module was associated with an indicator reaction used for EFM sampling. Aging-regulated modules were then identified by assessing the enrichment of aging-associated reactions, as indicated by the translation of aging-regulated genes into their corresponding metabolic reactions.

**Figure 6:**
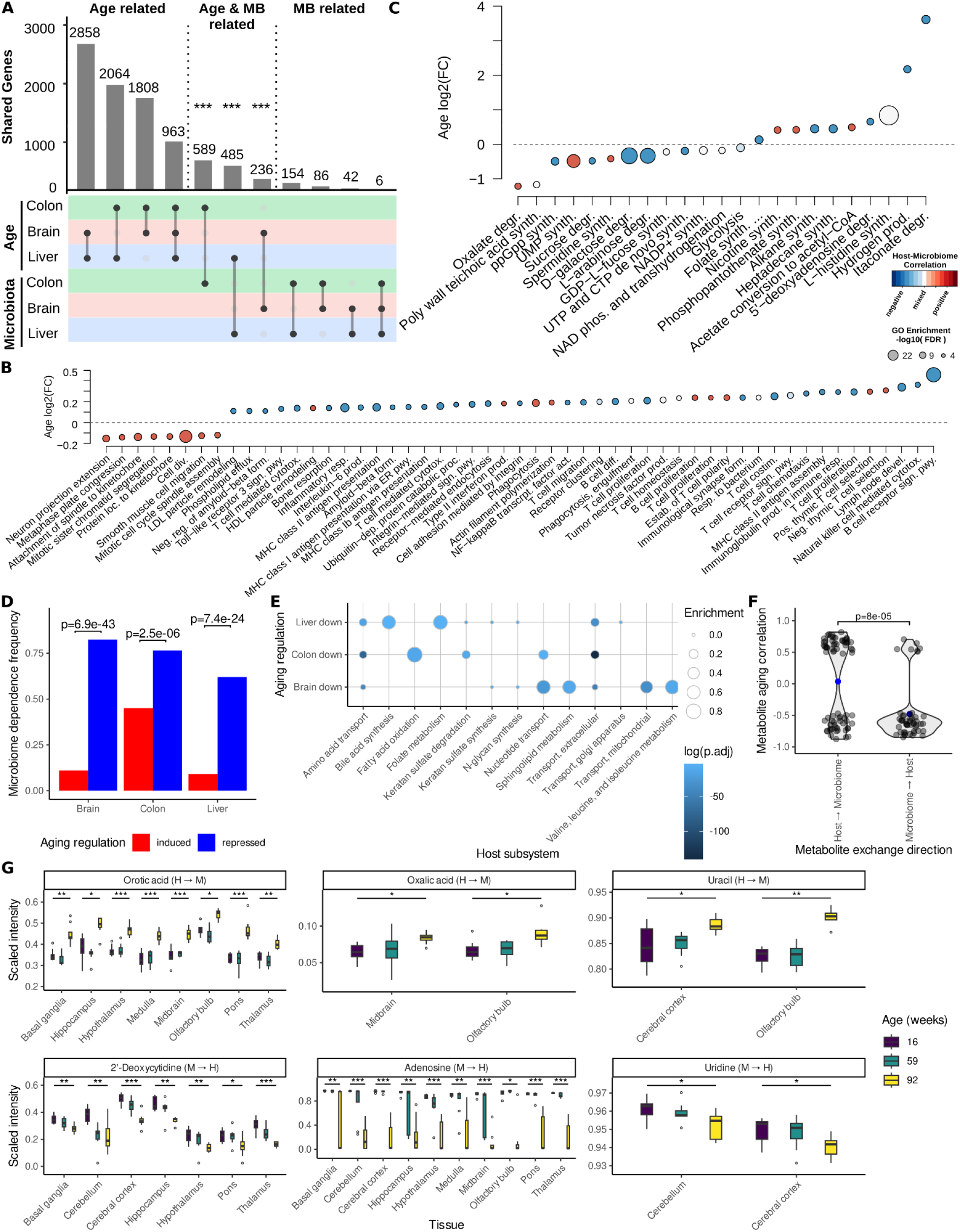
Aging-associated changes in host–microbiome interactions. **A** Overlap between aging-regulated and microbiome-regulated host genes. Asterisks indicate significant enrichment in the overlap according to a hypergeometric test. **B** Colon-specific gene expression changes with age in GO biological processes that were also correlated with microbiome metabolic functions. **C** Aging-dependent changes in microbiome functions that were correlated with host gene expression. **D** Frequency of microbiome dependence of aging-repressed and aging-induced metabolic modules across host tissues. The metabolic modules are assumed to be microbiome-dependent if they include at least one microbial reaction. The p-values indicate significance as determined by Fisher’s exact test. **E** Enrichment of the indicator reactions of aging-regulated metabolic modules among annotated metabolic host subsystems. The x-axis represents host metabolic subsystems, and the y-axis represents the corresponding aging-regulated gene sets. **F** Comparison of aging-associated changes in the brain concentrations of metabolites produced or consumed by the microbiome. Data were obtained from Feng et al. ^51^ (see Supplementary Table S6.4 for the complete dataset). **G** Aging-associated metabolome changes for selected model-predicted microbiota-produced and microbiota-consumed metabolites. Data were obtained from Feng et al. ^51^. Asterisks indicate the significance of differences: *, p ≤ 0.05; **, p ≤ 0.01; ***, p ≤ 0.001.

In the colon, liver, and brain, we identified significant enrichment in pathways for aging-induced (51, 88, and 99, respectively) and aging-repressed (2509, 1702, and 524, respectively) metabolic modules (Supplementary Tables S6.1–S6.3), which showed a much greater number of aging-repressed metabolic modules despite the lack of bias in the aging regulation of genes (Supplementary Tables S5.1–S5.3). Focusing on metabolic modules dependent on the microbiota (see Methods), we observed that aging-repressed modules were more frequently microbiome-dependent across all tissues (Fig. 6D). Further analysis of microbiota-dependent aging-regulated modules revealed a notable downregulation of colon metabolic modules related to amino acids, nucleotide uptake, and fatty acid oxidation, which are central to cellular homeostasis (Fig. 6E), along with a downregulation of metabolic modules involved in fatty acid oxidation. In the liver, we observed the greatest enrichment of downregulated microbiome-dependent metabolic modules in bile acid synthesis and folate metabolism, consistent with the increases in host-regulated bile acids and decreases in microbiome-regulated bile acids in the feces with age (Fig. 4F). In the brain, aging-regulated microbiome-dependent metabolic modules were enriched in nucleotide metabolism, sphingolipid metabolism, and branched-chain amino acid metabolism.

Given that we observed a particularly strong effect of aging-suppressed microbiome metabolic function on the brain (cf. Fig. 6D,E), we investigated the impact of aging on the microbiome-dependent metabolome in the brain by employing previously published brain metabolome data ^51^. To achieve this, we determined the correlations between metabolite concentrations and the age of the mice and separated the metabolites into two groups: those predicted by the metamodel to be provided by the microbiota to the brain and those predicted to be provided by the brain to the microbiota. Similar to our findings for the fecal metabolome data, metabolites provided by the host to the microbiota accumulated in the brain with age, whereas microbiota-produced metabolites were depleted (Fig. 6F). This involved nucleotide precursors, such as orotate and uracil, which accumulated in the host, whereas the intermediates and products of nucleotide salvage, such as adenosine, 2-deoxycytidine, and uridine, were depleted (Fig. 6G).

## Discussion

In this study, we performed a comprehensive model-based analysis of aging-associated alterations in the interactions between the host and its microbiota in mice. We acquired 181 MAGs using shotgun and long-read sequencing techniques, which were then converted into constraint-based metabolic networks. Our investigation into the correlations between host transcript levels and microbiome metabolic functions revealed extensive associations between the microbiome and host organs, even remote organs such as the brain. On the host side, many of these correlations were linked to immune processes, mitochondrial function, and chromatin modification, whereas on the microbiome side, they were linked to metabolites with known immune-modulatory functions. These metabolites included D-galactose (which has an immunomodulatory effect on the microbiome ^52^ and an effect on aging ^53^ in mice) and leucine (an important regulator of T-cell function ^54^). Moreover, we observed strong associations between host functions and microbial fermentation and nucleotide metabolism pathways, consistent with the central role of microbe-produced short-chain fatty acids and nucleotides in colonic energy balance ^55^ and intestinal barrier function ^56^, respectively.

One key component of our analysis was the reconstruction of metabolic metaorganism models for each mouse from transcriptomic and metagenomic data, which allowed us to propose the underlying pathways that might mediate associations between the host and the microbiota. These metamodels followed a similar setup to whole-body metabolic models previously reconstructed for humans ^57^. Reassuringly, these metamodels were able to recover many well-known metabolic microbiome–host interactions involving, for example, short-chain fatty acids, bile acids, and essential amino acids.

We found that host genes that were correlated with microbiome functions were highly enriched for model-predicted interactions between the host and the microbiota. A closer inspection of the reaction-level dependencies between the host and the microbiota via EFM analysis revealed that the host most often depended on central metabolic microbiome reactions. From an evolutionary perspective, a host’s dependence on central metabolic microbiome functions is plausible because they are likely present in many bacteria, reducing the host’s dependence on specific bacterial species and providing a larger pool of potential interaction partners ^58^. In addition, this dependence is consistent with prior observations of a much stronger conservation of microbiome functions compared with microbial species in the human gut microbiota across cohorts ^59^.

Another interesting aspect of the predicted microbiome–host exchanges is metabolites that the host can produce itself, such as nucleotides. The reasons for the existence of such exchanges could include advantages from a division of labor, as frequently observed within microbial communities ^60^, a reliance of the host on the microbiota as a metabolic backup system to increase phenotypic plasticity ^61^, or evolutionary addiction, whereby mutual dependencies develop due to the constant exposure of the host to microbially produced metabolites ^62^.

Having characterized the potential interactions between the host and the microbiome in-depth, we next investigated how they changed in the context of aging. We observed significant decreases in microbiome metabolic activity and growth, including in microbial production of the short-chain fatty acid butyrate, consistent with previous observations of reduced butyrate levels in the serum of aged mice ^41^ and humans ^63^. In contrast, metabolic modeling predicted increased production of the proinflammatory metabolite succinate ^64,65^, which has been previously shown to be an indicator for a dysbiotic gut environment ^49^ and is consistent with the deregulation of succinate metabolism and succinate-dependent regulatory pathways involving the aging-regulator SIRT5 on the host side ^66^. Moreover, our observation of reduced microbial growth with age could provide an additional mechanism mediating the increased likelihood of constipation ^67^ and longer colonic transit times ^68^ during aging since the latter predominantly depend on microbial growth rates ^69^.

Our analysis of predicted ecological interactions in the aging gut microbiome indicated that these changes are likely driven by an increase in competitive metabolic interactions within the gut microbiota with age alongside a decrease in cross-feeding interactions and, thereby, less efficient utilization of dietary resources. The reduced metabolic activity of the aging microbiome was also reflected in fecal metabolomics data, in which most identified metabolic features showed a reduction with age. In particular, host-regulated bile acids increased with age, while microbiome-regulated bile acids decreased. Moreover, we found that anti-inflammatory metabolites such as valine, betaine ^70,71^ and 3-hydroxykynurenine ^72^ decreased while pro-inflammatory metabolites such as D-galactose ^52^ increased. On the host side, we found appreciable conservation of aging-associated changes across tissues, including the induction of inflammatory processes and suppression of cellular replication, consistent with the conserved signature of aging that we identified previously ^3^. Our results for the individual organs were also consistent with prior observations, including altered gut motility (linked to enteric neurons) ^73,74^, reduced colonic barrier function ^14^, reduced proliferative potential ^75^, and altered mitochondrial morphology and reduced mitochondrial DNA in the liver of aged mice ^76^.

Considering aging-associated changes in host–microbiome interactions, we found that aging-regulated host genes were highly enriched for genes that were correlated with microbiome functions. In particular, downregulated metabolic modules in the host depended on the microbiota. These suppressed microbiome-dependent metabolic modules were particularly enriched among processes centrally involved in cellular homeostasis and proliferation, such as amino acid and nucleotide uptake. Importantly, these trends were also evident in correlation analyses in which host homeostatic processes were positively associated with microbial metabolic processes, whereas immune-associated host pathways were negatively associated with these processes. This demonstrates that the aging-associated reduction in microbial metabolism is negatively associated with host homeostatic processes and positively associated with inflammation. The observed loss of host–microbiome interactions across all organs indicates that the microbiome might contribute to crucial aspects of the systemic aging process, such as metabolic decline ^77,78^ and the loss of cellular proliferation, along with stem cell exhaustion ^2^.

We identified microbially-produced nucleotides as a key metabolic exchange in host–microbiome interactions. According to the model predictions, this interaction involved the host’s provision of nucleotide precursors, such as orotate, and nucleotide degradation products, such as uracil, to the microbiota, while the microbiota provided nucleotides and nucleotide salvage products in return. This interaction is further supported by the widespread correlations between host gene expression and microbial processes involved in nucleotide metabolism across all tissues examined. Although the host can synthesize all nucleotides *de novo* and obtain nucleotides from its diet, recent studies have indicated that the microbiota are a key source of nucleotides, particularly in the colon ^56,79^. On the microbiome side, *Escherichia coli* and *Bacteroides spp.* have been reported to excrete ATP during growth ^80,81^. Moreover, microbially-sourced nucleotides were identified as important contributors to intestinal barrier function ^56^. Furthermore, the host’s sensing of microbially-produced ATP by purinergic receptors in the colon is an important modulator of germinal center-mediated immune reactions toward the microbiota ^79^. In this context, our observation of an aging-associated loss of host–microbiome nucleotide co-metabolism may be critical in key aspects of host aging, including the documented decline in intestinal barrier function ^56^ associated with various age-related diseases ^82–84^, the decreased cellular proliferative capacity in the intestine and systemically ^85,86^, and the impaired mitochondrial function with age ^87^. According to the model predictions and brain metabolome data, the microbiome may play a key role in nucleotide salvage in the brain, which is essential for DNA damage repair and cellular homeostasis ^88^. In addition, an association between neurodegeneration and microbial pyrimidine metabolism was observed in an experimental mouse model of Alzheimer’s disease ^89^.

In summary, we comprehensively characterized microbiome–host interactions in mice and identified pronounced aging-associated changes in these interactions primarily driven by a loss of microbial metabolic activity. Although our study is limited by its reliance on modeling, we predicted many known microbiome–host interactions and attempted to corroborate model-based predictions through complementary independent analyses. Notably, although the metabolic metamodels identified that specific metabolites are exchanged, the promiscuity of transporters and the underlying modeling procedure might mean that compounds with closely related molecular structures are exchanged *in vivo*. Moreover, since our mouse cohort was exclusively comprised of males to avoid confounding due to sex differences, we could not detect sex-specific changes; future studies should apply our modeling approach to investigate this. Additionally, we based our analysis only on chronological age and did not assess more fine-grained markers of biological age, such as frailty, loss of motor function, or cognitive function impairment. Finally, our identification of a loss of microbiome metabolic activity indicates a potentially crucial aging-associated change that could contribute to many aging-associated pathologies in the host. Therefore, microbiome metabolic activity could be a target for future microbiome-based therapies. Our modeling approach could be a key ingredient in designing such interventions to guide the development of therapies counteracting microbiome-driven aspects of aging.

## Methods

### Mouse strain

The mice used in this study were an in-house strain derived from the C57BL/6J strain (The Jackson Laboratory, Bar Harbor, ME, USA). These C57BL/6J/Ukj mice lack two common mutations found in the C57BL/6J strain: the DIP686 mutation in the crumbs family member 1 (*Crb1*) gene, which is vital for eyesight in aging mice, and a mutation in the nicotinamide nucleotide transhydrogenase (*Nnt*) gene, which encodes mitochondrial NAD(P) transhydrogenase, protecting against oxidative stress. Preserving both these genes is advantageous for metabolic and aging studies in mice.

### Animal handling

Male C57BL/6J/Ukj mice were bred in the Central Experimental Animal Facility at Jena University Hospital (Jena, Germany). The mice were housed at 22 ± 2°C with a 14/10 h day/night cycle and a relative humidity of 55% ± 10%. They were co-housed according to their birth cohort (similar ages) in standard cages (GM500, Type III; Tecniplast Deutschland GmbH, Hohenpeißenberg, Germany), and a maximum of two mice from the same cage were used for experiments. The mice had unlimited access to water and food (mouse V1534-300, ssniff Spezialdiäten GmbH, Soest, Germany). Next-generation RNA sequencing of host tissues and metagenomics of fecal samples were conducted in 52 mice of different ages spanning the mouse’s adult lifespan (2–3 [mean = 2.5] months, 9–10 [mean = 9.8] months, 15–17 [mean = 15.9] months, 24–25 [mean = 24.8] months, and 28–31 [mean 29.1] months). For simplicity, the five age groups are referred to as 2 (*n* = 10), 9 (*n* = 10), 15 (*n* = 10), 24 (*n* = 10), and 30 (*n* = 12) months throughout the manuscript (see Supplementary Table S1.1).

An independent mouse cohort was used for the metabolomics analysis of fecal samples; this cohort comprised 83 male mice in five age groups: 3 (*n* = 16), 9 (*n* = 16), 15 (*n* = 16), 24 (*n* = 17), and 28 (*n* = 18) months (see Supplementary Table S4.11). These mice were bred and housed in the same mouse facility under the same conditions.

### Sample collection

The mice were sacrificed by cervical dislocation in three cohorts (randomized by age) on three consecutive mornings. The left hemisphere of the brain was prepped on ice, transferred to liquid nitrogen for storage, and used later for RNA extraction. Feces were collected from the colon by squeezing the colon contents toward the distal end and snap-freezing one pellet in liquid nitrogen; the pellets were used later for metagenomic sequencing (for the first cohort) or metabolite measurement by hydrophilic interaction liquid chromatography (HILIC) ultrahigh-performance liquid chromatography (UHPLC)-tandem mass spectrometry (MS/MS) (for the second cohort). The colons were rinsed with sterile PBS and cut longitudinally; a piece measuring the length of one-eighth of the left half of the mid colon was frozen in liquid nitrogen for later use for RNA extraction. A piece with a length of approximately 1 cm length was cut from the end of the left lateral lobe of the liver and snap-frozen in liquid nitrogen for later use for RNA extraction.

All studies were performed in strict compliance with the recommendations of the European Commission for the protection of animals used for scientific purposes and with the approval of the local government (Thüringer Landesamt für Verbraucherschutz, Germany; license: 02-024/15; TWZ-000-2017). Experiments were performed according to the ARRIVE guidelines

### Metagenomic sequencing

Microbial DNA was extracted from colon contents with the DNeasy PowerSoil Kit (Qiagen, Hilden, Germany) following the manufacturer’s protocol. Next, the DNA was prepared at the Max Planck Institute for Evolutionary Biology (Plön, Germany) with the Illumina NexteraXT Library Kit. All 52 samples were pooled and sequenced for 2 × 150 cycles in paired-end mode on all four lanes of an Illumina NextSeq 500 machine. Demultiplexing was performed with one mismatch allowed in barcodes. The raw read data were merged sample-wise and subjected to quality control for adaptor contamination and base call qualities. Adaptor sequences with an overlap of ≥3 bp and base calls with a Phred+33 quality score of <30 were trimmed from the 3′ ends of reads using Cutadapt (version 1.12). Illumina’s Nextera transposon sequence and the reverse complement of TruSeq primer sequences were used as adaptor sequences.

Subsequently, reads were subjected to quality control using Prinseq lite (version 0.20.4) with a sliding window approach that applied a step size of 5 bp, a window size of 10 bp, a mean base quality of <30, and a minimum-length filter that discarded any reads shorter than 50 bp after all other quality control steps. To filter out host sequences, the remaining sequences were mapped to the mouse reference genome (GRCm38.99) with Bowtie (version 2.2.5). The remaining unmapped reads were then used for MAG assembly.

No significant differences were detected in the total microbial read depth or host contamination between age groups (Kruskal–Wallis test with post hoc Dunn’s test and Benjamini–Hochberg multiple testing correction conducted with the DunnTest function in the DescTools R package [version 0.99.50]; see Supplementary Figs. 1C and 1D).

Long-read sequencing was performed at the NGS core facility of the FLI Leibniz Institute on Aging (Jena, Germany). The DNA quality was assessed with an Agilent Bioanalyzer 2100 with a DNA 12000 Kit (Agilent Technologies, Santa Clara, CA, United States) and quantified with an Invitrogen Quant-iT PicoGreen dsDNA Assay (Thermo Fisher Scientific, Waltham, MA, United States). The sequencing library was prepared according to the Pacific Biosystems’ manual “Procedure & Checklist-20 kb Template PreparationUsingBluePippin Size-SelectionSystem” (version 10, Jan. 2018) with the SMRTbell Template Prep Kit 1.0 (Pacific Biosciences, Menlo Park, CA, United States). Specifically, DNA from age-matched samples was pooled, fragmented (75 kb) by a Megaruptor (Diagenode, Denville, NJ, United States), and size-selected for >6 kbp fragments with a BluePippin and 0.75% Gel Cassette (program: 0.75% DF Marker S1 High-Pass 6–10 kb vs3; Sage Science, Beverly, MA, United States). Each pool was loaded onto a SMRTcell and sequenced on a Pacific Biosystems RSII machine with DNA-Sequencing Kit 4.0 v2, MagBeadBuffer Kit v2, MagBead Binding Buffer Kit v2, and DNA Polymerase Kit P6v2. The sequence output of this run had an average read length of 7.8–9.7 kb with a minimum yield of 750 kbp per pool/SMRTcell. The raw read data were subjected to quality control, processed into circular consensus sequences and subreads, and exported as FASTQ files via the SMRTportal (provided by Pacific Biosciences).

### MAG assembly and annotation

MAGs were constructed as follows (outlined in Supplementary Fig. 1B). Pacific Biosystems circular consensus sequences and subreads were used as is, while Illumina shotgun reads were filtered for low read quality, adapters, and host contamination (see the Metagenomic Sequencing section). A full cohort assembly was done in metaSPAdes (SPAdes version 3.13.1) in hybrid mode with *k*-mer sizes of 21, 33, 55, and 77. Concatenated, quality-controlled, forward and reverse Illumina short read files of all samples were used as input. Additionally, the assembly software was informed with the eight Pacific Biosystems long read banks (hybrid mode) in the form of filtered subreads and circular consensus sequences.

The resulting scaffolds were filtered for a minimum length of 1000 bp and coverage ≥7.7815. The cutoffs were determined by scatter plotting coverage versus length, as described by ^91^. The quality-controlled metagenomic reads were mapped back to the filtered scaffolds with Bowtie (version 2.2.5); the insert size was 0–1000 bp in the very sensitive, non-deterministic, “fr” stranded mode with end-to-end alignment. Non-unique mappings and unaligned reads were discarded. The scaffold coverage depth was determined with the jgi_summarize_bam_contig_depths script from MetaBAT (version 2.12.1). This coverage depth information was then used to sort the remaining scaffolds into bins, each representing single bacterial genomes, with the binning tools MetaBAT (version 2.12.1), CONCOCT (version 1.1.0), and MaxBin (version 2.2.4). For CONCOCT, the scaffolds were broken up into 10 kbp chunks. Bin refinement was conducted with the combined results of all three binners (252 bins) with DASTool (version 1.1.2); subsequently, quality metrics were calculated by CheckM (version 1.1.2). Bins with a quality estimate of >80% and a contamination estimate of <10% were considered for further analysis and are henceforth referred to as MAGs. The 181 final MAGs were taxonomically annotated with GTDB-Tk (version 2.1.1) and database version r214. The tRNA genes were characterized using tRNAscan-SE (version 2.0.9) and in-house Bash scripts. The 16S rRNA genes were detected by barrnap (version 0.9) in “kingdom bacteria” mode. A phylogenetic tree of the 181 MAGs (Fig. 1A) was created from a multiple sequence alignment created by GTDB-Tk (align/gtdbtk.bac120.user_msa.fasta.gz) with the European Bioinformatics Institute’s online Simple Phylogeny tool (ClustalW version 2.1) and visualized with R statistical software. The complete characterization of the MAGs is provided in Supplementary Table S1.2. For association with age, MAG abundances were calculated from the scaffold coverage depths described above, normalized by sample, and then correlated with age in linear models for each MAG across all samples. The *p*-values were corrected for multiple testing with the Benjamini–Hochberg FDR method. Significant age-associated MAGs (FDR-adjusted *p*-value ≤0.05) were plotted (Fig. 4A).

### Microbiome metabolic model construction

Metabolic models were constructed for each of the 181 murine gut bacteria inferred from our MAGs in samples from the 52 mice. The reconstruction was performed in gapseq (version 1.2) with default settings (git commit: 159ad378; sequence DB md5sum: bf8ba98) ^31^. The nutritional input for the computational models was designed according to the fortified rat and mouse diet (V1534-300; ssniff Spezialdiäten GmbH). The diet was reconstructed according to the vendor’s information on its molecular constituents translated into the corresponding metabolites in the models (following the protocol described in Marinos et al. ^92^). We assumed an average daily uptake of 3.5 g of food based on reference values^93^. This amount was used to transform the percentages into grams and then millimoles (millimoles/day). Limited information was reported on fiber in the mouse diet; therefore, their values were imputed from the consumed quantities of cereal and grain products of a human German cohort ^23^. Because the simulations depicted the intestinal setting, the absorption in the small intestine was considered when calculating the dietary input (see Supplementary Tables S1.3-S1.11 for the respective calculations and references).

### Comparing metabolic models via principal component analysis

Metabolic models were used to create an incidence matrix of all available reactions per model. The incidence matrix was normalized sample wise in order to weigh the importance of reactions by the size of the model. This means larger models with more reactions had lower weight per reaction compared to smaller models containing less reactions. Dissimilarity between bacterial metabolic models was calculated from this incidence matrix via Horn–Morisita index as implemented in the function vegdist from R-package vegan v2.6-4. Principal Component Analysis was performed on these dissimilarity indices via function prcomp from R-package stats v4.3.2. Metadata information of each model, namely count of tRNAs, completeness, contamination, GC-content, genome size, model size and model gaps, was fitted to the principal coordinates with the function envfit (vegan v2.6-4). The first two principal components were plotted together with the metadata vectors in Fig. 1B. Explained variance for model fitting of each metadata variable separately against the first two principal components was reported as R^2^ values in the text.

### Estimation of functional capacity of microbiomes

Flux variability analysis (FVA) was used to calculate all possible flux ranges for each reaction in each metabolic model constrained by the ssniff diet. The biomass production was set as the objective function to obtain flux ranges that satisfied maximal model growth (99% percent of maximal growth used as cut-off). A binary incidence matrix was constructed with bacteria as columns and biochemical reactions as rows. All reactions that carried a non-zero (>10^−6^) flux were assigned “1”. This matrix was first normalized to the number of active reactions in each bacterium and then multiplied by each mouse’s bacterial abundances to obtain a reaction abundance for the microbiome of each mouse. Finally, the reaction abundance table was normalized by the sum of each mouse’s reaction scores, making the resulting relative reaction abundances comparable between mice.

### Growth rate prediction from metagenomic data

To further validate the model growth rates, CoPTR software ^94^ was used to estimate growth rates from the MAGs in each sample. This software uses the peak-to-trough ratio (PTR) (i.e., the ratio of sequencing coverage near the replication origin and the replication terminus) to estimate the growth of a MAG in a sample ^95^. We first indexed the MAGs with the command “coptr index--bt2-threads”. Next, using this index, we mapped our quality-controlled metagenomic reads against our 181 MAGs with the command “coptr map--threads 4--paired”. Then, read positions were extracted with the command “coptr extract,” and the PTR was estimated with the command “coptr estimate.” Default parameters were used for all commands except “coptr index” and “coptr map” for which the number of threads was specified. Additionally, “--paired” was set for the “coptr map” command to inform the software about the use of paired-end reads. Community growth was determined for each mouse’s microbiome community by calculating the median growth rate across all MAGs in its sample.

### Inferring microbial niche strategies via the universal adaptive strategies theory

Microbial life history traits, defined by the universal adaptive strategies theory (UAST) framework ^44,96^, were predicted (similar to a previous study ^45^). Microbial traits were inferred by gapseq (version 1.2; sequence DB md5sum: bf8ba98) ^31^ using MetaCyc pathways ^33^, Bakta v1.8 ^97^, abricate (version 1.0; https://github.com/tseemann/abricate) with the virulence factor database ^98^, and gRodon (version 2.3) ^99^. The competitive strategy was defined by the following traits: genome length (high quantities), antibiotic biosynthesis pathways (high), siderophore biosynthesis pathways (high), and catabolic pathways (high). The stress toleration strategy was defined by rRNA genes (low quantities), biofilm genes (high quantities), and auxotrophies (high quantities). The ruderal strategy was defined by catabolic pathways (low quantities), rRNA genes (high quantities), and codon usage bias (high quantities). Given the distribution of each trait among all species, a species’ trait belonging to the 0.75 (traits with high quantities) or 0.25 (traits with low quantities) quantile was considered to contribute to a strategy. The number of contributing traits was summed for each strategy, and the strategy with the highest number of contributing traits was considered a species’ final life history strategy. In the case of multiple strategies with the same highest number of contributing traits, multiple strategies were assumed to be relevant. In addition to the 0.75 and 0.25 quartiles, different percentiles (0.7/0.3 and 0.8/0.2) were tested to assess stability.

The niche strategies of all species from one microbial community (per mouse) were summed, with a down-weighting of the impact of species with two equally likely strategies by 0.5 and a down-weighting of the impact of species with three equally likely strategies by 0.333. The niche strategy abundances were normalized by sample and multiplied by 100 to obtain percentages. Differences between age groups were tested for each of the three possible niche strategies using Kruskal–Wallis tests with pairwise post hoc Dunn’s tests and Benjamini–Hochberg FDR correction (using the DunnTest function of the R DescTools package [version 0.99.50]) and plotted (Supplementary Fig. 4B).

### Microbiome Community Modeling

The microbial communities were modeled using FBA to study the association between the microbiome metabolic network and the age of the mouse hosts. FBA is a mathematical approach for studying metabolic networks built from all known reactions in an organism. By estimating the flow of metabolites in the network, FBA allows the prediction of the growth rate of the organism and the fluxes of all metabolites. This is carried out under the evolutionary assumption that the preferable path maximizes the biomass compounds. For prediction of microbial community fluxes, we used community FBA ^23^, a variant of FBA working on the community level. To this end, the metabolic networks of different microbial species within the community were connected in a common compartment for metabolic exchange within the community and with the environment (the host’s intestinal tract). A community-level biomass reaction was introduced draining the biomass of individual species according to their abundance as inferred from metagenomic data. Additionally, we introduced coupling constraints to prevent excessive flux through individual microbes’ reactions’ without concomitant growth (coupling parameters c=400, u=0.01). For predicting maximal growth, we optimized the community-level biomass reaction while subtracting the total sum of fluxes across all reactions multiplied with a factor of 10^-6^ to obtain a parsimonious solution. No feasible solution could be obtained for two microbiome communities from age group 30, leaving 50 communities for downstream analysis (10 for each age group).

This analysis identified three types of reaction fluxes. Exchange fluxes of metabolites exported and taken up from outside the community were considered metabolites that may be exchanged with the host (Fig. 4C). Metabolites shared between different microbes were represented by reaction fluxes that were exchanged among members of the microbial community (Supplementary Fig. 4A). Finally, the reaction fluxes within each respective bacterial model were considered as internal reaction fluxes (Fig. 4B).

All three types of reaction fluxes were normalized to the community growth rate and analyzed separately. Absolute flux values were correlated with age using Spearman’s rank correlation coefficient (using the cor.test function in the R package stats [version 4.3.2]) separately for each reaction flux. The *p*-values were corrected for multiple testing with the Benjamini–Hochberg FDR method, and only results with FDR-adjusted *p*-values of ≤0.05 were plotted (Fig. 4B, C, and Supplementary Fig. 4A).

Significant age-correlated internal reaction fluxes were enriched for MetaCyc pathways, stratified by positive and negative correlations, with an overrepresentation test implemented in the enricher function of the R package clusterProfiler (version 4.8.3). The pathways with an enrichment *p*-value of ≤0.05 and at least three enriched features were reported (Fig. 4B; Supplementary Tables S4.2–S4.4).

The ecological relationships for each pair of bacteria across all species were predicted. To this end, the growth achieved as a single bacterium was compared with that achieved when each bacterium was co-grown with another. The relationships were characterized using the ecological relationships described by Zélé et al. (Figure 1 in Ref. ^100^). as a reference. Growth was estimated by FBA for single growth and community FBA for combined growth. To achieve this, we used the R packages sybil ^101^ and MicrobiomeGS2 (www.github.com/Waschina/MicrobiomeGS2) and the linear programming solver IBM ILOG CPLEX 22.10. The MicrobiomeGS2 simulations followed the principles described previously ^27^. The six types of ecological relationships and their frequencies among each microbial community were inferred with the R EcoGS package (https://github.com/maringos/EcoGS).

To obtain relative frequencies, the abundance of ecological relations was normalized by sample to a sum of 1. Next, a linear model analysis of each ecological interaction type with age was conducted and *p*-values were adjusted for multiple testing using the Benjamini–Hochberg FDR method.

### Transcriptome Sequencing

RNA was extracted from tissue samples of the liver, colon, and left brain hemisphere using the phenol-chloroform extraction method with 1 mL of Qiazol Lysis Reagent (Qiagen, Hilden, Germany), as previously described in Ederer *et al.* ^102^. Next, the RNA was reverse-transcribed into Illumina shotgun sequencing libraries with TruSeq RNA stranded kit and polyA enrichment following the manufacturer’s protocol (Illumina, San Diego, CA) at the Competence Centre for Genomic Analysis (Kiel, Germany) and sequenced for 2 × 75 cycles (2 × 100 for colon) in paired-end mode with ∼13 samples per lane on an Illumina HiSeq4000 machine. Demultiplexing was conducted with zero mismatches allowed in the barcodes. Illumina TruSeq adapter sequences were trimmed from the forward and reverse reads with Cutadapt (version 1.12) with a minimum sequence overlap of 3 bp and no more than 10% mismatches; they were also filtered for a minimum read length of 20 bp and trimmed for 3’-end quality using a Phred score of ≥30. Additional quality filters were applied with Prinseq lite (version 0.20.4), allowing at most eight unknown nucleotides (“N”) per read and requiring an overall mean Phred score (read quality) of ≥15; the reads were also trimmed for 5’-end quality using a Phred score of ≥12.

The filtered reads were mapped against the *Mus musculus* reference genome (GRCm38.99) in Hisat2 (version 2.1.0) with the RNA strandedness set to FR, applying non-deterministic random seeds and suppressing mixed alignments of read pairs. Only primary alignments for each read were kept, via a filtering step (-F 256) in Samtools (version 1.9). Gene counts were extracted in HTSeq count (version 0.6.1) with reverse-stranded information in union mode.

### Differential gene expression analysis

Differentially expressed genes were identified using the R package DESeq2 (version 1.40.2) ^103^.The samples were stratified by organ (colon, liver, and brain) and then analyzed with a design formula that accounted for the age (numeric, centered, and scaled) and the sequencing batch, if applicable (liver and brain). Genes differentially expressed according to age were reported, controlling for independent filtering at 0.05 using DESeq2 (function: results, variable: alpha). Genes with an adjusted *p*-value of ≤0.05 were considered significantly differentially expressed. Differentially expressed genes were stratified by their positive or negative association with age and annotated separately with GO biological processes via enrichment analysis using the enricher function of the R clusterProfiler package (version 4.8.3). For plotting (Fig. 5), the GO terms were filtered using an FDR-adjusted *p*-value cutoff of 10^−4^ for the colon and 10^−6^ for the liver and brain; redundant higher-level GO terms were then removed, as described in ref. ^104^.

Variance-stabilizing transformed gene abundance data were exported for downstream use with host–microbiome partial correlations using DESeq2 (function “getVarianceStabilizedData” with parameter blind = FALSE).

All samples were jointly loaded and processed with a design formula accounting for the sequencing batch and age group. The results were stratified by organ and extracted from the DESeq results object for each of the 10 possible pairwise age group comparisons.

### Host–Microbiome Partial Correlations

The transcriptomic data was normalized separately for each organ (colon, liver, and brain) using variance-stabilizing transformation informed with age and sequencing batch (blind = FALSE) implemented in the R package DESeq2 (version 1.40.2) ^103^. A near-zero variance filter was also applied using the nearZeroVar function of the R package caret (version 6.0-94). The active reactions of each mouse’s microbiome community were predicted as described in “Estimation of Functional Capacity of Microbiomes”. The host transcript abundances were correlated pairwise with microbiome active reactions (each transcript with each reaction), correcting for age and sequencing batch (only for the liver and brain), with Spearman’s partial correlations (implemented in the R package ppcor [version 1.0] ^105^). To balance stringent false discovery cutoffs with reasonable result counts, strong correlations with a Benjamini–Hochberg FDR-corrected ^106^ *p*-value of ≤0.1 and Spearman’s *ρ* of ≥0.55 were considered for downstream analysis. The strong correlations were stratified into either positive or negative correlations according to their correlation values and then annotated with GO biological processes ^32^ (host transcripts) or MetaCyc Pathways ^33^ (microbiome reactions) using hypergeometric overrepresentation tests with the phyper function of the R stats package (version 4.3.2; *x* = *“correlated features enriched for the term” − 1*, *m* = *“the total of all correlated features,” n* = *“all features” – “correlated features,”* and *k* = *“the total of the features in the term”*). Enriched terms (pathways/processes) with at least three features and an FDR-corrected overrepresentation *p*-value of ≤0.05 were reported (Supplementary Tables S2.1–S2.3; Fig. 2). The negative decadic logarithm of the overrepresentation FDR *p*-values was calculated and reported as-is for positive correlations and multiplied by −1 for negative correlations. Only process pairs that were associated with at least two other pathways were plotted (Fig. 2) and filtered to highlight the most significant enrichments with FDR *p*-value cutoffs of ≤1·10^−10^ for the colon (Fig. 2B), ≤1·10^−4^ for the liver (Fig. 2D), and ≤1·10^−3^ for the brain (Fig. 2F).

Correlated feature pairs were obtained for the colon (*n* = 12,732), liver (*n* = 3,425), and brain (*n* = 2,499). They consisted of *n* unique features for the colon (microbiome, *n* = 1,606; host, *n* = 2,815), liver (microbiome, *n* = 1,359; host, *n* = 1,277), and brain (microbiome, *n* = 1,236; host, *n* = 926). After enrichment, we obtained *n* process pairs for the colon (*n* = 1,377), liver (*n* = 283), and brain (*n* = 167), as shown in Fig. 2 and Supplementary Tables S2.1–S2.3.

For a broader overview of host–microbiome associations, the GO biological processes were grouped by their higher-ranking level 2 GO biological process, and the MetaCyc pathways were grouped by their respective highest-level super-pathways (see Supplementary Tables S2.5 and S2.6 for the process and pathway groups). The level 2 GO biological process groups were cellular process, metabolic process, biological regulation, localization, developmental process, response to stimulus, immune system process, multicellular organismal process, viral process, reproduction, homeostatic process, and growth. The MetaCyc microbial super-pathways were lipids, carbohydrates, utilization, energy metabolism, nucleotides, secondary metabolites, amino acids, other, signaling, carboxylates, cofactors, carriers, metabolic regulators, c1 compounds, electron transfer, noncarbon nutrients, cell structure, biosynthesis, detoxification, interconversion, glycans, tRNA, and bioluminescence. The −log_10_(FDR-corrected *p*-values) were summed for each level 2 GO and MetaCyc superpathway pair. The values are plotted in Fig. 2A, C, E as well as Supplementary Fig. 2 and listed in Supplementary Table S2.4.

The pairwise correlations between all host features and all microbiome features were repeated, stratified by organ and age group. Thus, the ratio of significant, strong host–microbiome correlations to all tested correlation pairs was obtained separately for each organ and age group. These ratios were compared using Pearson’s Chi-squared test with Yates’ continuity correction and Bonferroni’s multiple testing correction to identify significant differences between consecutive age groups (Supplementary Fig. 6A–E).

The overlaps of aging-regulated and microbiome-associated transcripts between the three studied organs were determined to identify shared aging-regulated and microbiome-regulated host transcripts. The overlap between microbiome-associated and aging-associated transcripts was statistically evaluated using hypergeometric overrepresentation tests separately for each organ. The numbers of shared transcripts were plotted (Fig. 6A) for each possible combination for the colon (age-associated, *n* = 4,715; microbiome-associated, *n* = 2,326; shared, *n* = 527; hypergeometric *p*-value = 6.9 × 10^−21^), brain (age-associated, *n* = 6,505; microbiome-associated, *n* = 888; shared, *n* = 260; hypergeometric *p*-value = 1.8 × 10^−11^), and liver (age-associated, *n* = 8,285; microbiome-associated, *n* = 1,265; shared, *n* = 500; hypergeometric *p*-value = 1.6 × 10^−9^).

### Non-targeted Metabolomics using HILIC UHPLC-MS/MS

Fecal pellets (∼40 mg) were weighed in sterile ceramic bead tubes (NucleoSpin Bead Tubes; Macherey-Nagel, Dueren, Germany) and extracted with 1 mL of chilled methanol (−20 °C; LiChrosolv, Supelco; Merck KGaA, Darmstadt, Germany). Fecal matter was homogenized and extracted with a Precellys Evolution Homogenizer (Bertin Corp., Rockville, MD, USA; 4,500 rpm, three 40-second cycles with a two-second pause). The samples were centrifuged for 10 minutes at 21,000 ×g and 4°C, and the supernatant was transferred into sterile tubes until analysis. Next, 100 µL of fecal methanolic extract was evaporated at 40°C with a SpeedVac concentrator (Savant SPD121P; Thermo Fisher Scientific, Waltham, MA, USA) and reconstituted in 75% acetonitrile (ACN; LiChrosolv, hypergrade for LC–MS; Merck KGaA), spiked with L-leucine-5,5,5-d_3_ at 5 mg/L (99 atom % D; Merck KGaA), which was prepared in a methanol and water solution (50:50).

The samples were analyzed by using a UHPLC system (Acquity; Waters, Eschborn, Germany) coupled to a quadrupole time-of-flight (TOF) mass spectrometer (maXis; Bruker Daltonics, Bremen, Germany), as described previously ^107^. Mass spectra were acquired at electrospray ionization of positive and negative modes (+/−). Spectrometric data were acquired in line and profile mode with an acquisition rate of 5 Hz from 50 to 1500 Da. Fragmentation experiments were set to the data-dependent mode (MS/MS [Auto]), where the three most intense ions were fragmented within one scan when a count reached over 2000. Ions were excluded after acquiring three MS/MS and reconsidered for fragmentation after six seconds. The collision energy was set to 20 eV for both modes with an isolation width of 8 Da. The electrospray ionization source parameters were as follows: capillary voltage of 4500 V for (+) and 4000 V for (−), end plate offset of (+/−) 500V, nebulizer gas of 2 bar, dry gas of 10 L/min, and dry heater of 200°C. Before measurements, the MS was calibrated using the ESI-L Low Concentration Tuning Mix (Agilent, Santa Clara, CA, USA). The ESI-L Low Concentration Tuning Mix (diluted 1:4 [v/v] with 75% ACN) was injected in the first 0.3 min of each run by a switching valve for internal recalibration by post-processing software.

Ammonium acetate (NH_4_Ac; LiChropur eluent additive for LC–MS; Merck KGaA) at 0.5 mol/L was adjusted to pH 4.6 with glacial acetic acid (Honeywell; Fluka, Seelze, Germany). Milli-Q water was obtained from a Milli-Q Integral Water Purification System (Billerica, MA, USA). Polar metabolites were separated by HILIC by using an iHILIC-Fusion UHPLC column SS (100 × 2.1 mm, 1.8 µm, 100 Å; HILICON AB, Umea, Sweden). The eluent compositions were as follows: Eluent A consisted of 5 mmol/L NH_4_Ac (pH 4.6) in 95% ACN (pH 4.6), and eluent B consisted of 25 mmol/L NH_4_Ac (pH 4.6) in 30% ACN. We started with 0.1% B, keeping it constant for two minutes, then increased B to 99.9% over 7.5 minutes. The condition of 99.9% B was kept for two minutes and reversed to 0.1% B within 0.1 minutes, held for 0.1 minutes. The run was completed after 12.1 minutes, and the column was equilibrated for five minutes before the next injection. The flow rate was set to 0.5 mL/min, the column temperature to 40°C, and the sample manager was cooled to 4°C; 5 µL of the sample was injected into the column (partial loop). The weak and strong washes consisted of 95% and 10% ACN, respectively.

### Metabolite Identification and Metabolomic Data Processing

The raw LC–MS data were post-processed in GeneData Expressionist Refiner MS (version 13.5.4; GeneData GmbH, Basel, Switzerland), including chemical noise subtraction, internal calibration, chromatographic peak picking, chromatogram isotope clustering, valid feature filter (cut-off of 100 [+] or 2000 [−] maximum intensity and presence of features in at least 20% of samples for [+/−]), retention time range restriction (0.4–10.7 minutes), annotation of known peaks (mass-to-charge tolerance of 0.01–0.005 Da and retention time tolerance of 0.1), and MS/MS consolidation and export to merged MASCOT generic files (MGFs). Data processing resulted in a matrix containing features with mass-to-charge ratios (*m*/*z*), retention times, and observed maximum intensities for each sample. The data were normalized to the weighed-in wet fecal weight and a maximum intensity of 5 mg/L L-leucine-5,5,5-d_3_.

The merged MGF files were used to search a spectral library using MSPepSearch (0.01 Da mass tolerance for precursor and fragment searches). Experimental and *in silico* spectral libraries were downloaded from MassBank of North America (https://mona.fiehnlab.ucdavis.edu/). Identification was performed by matching experimental MS/MS spectra against MS/MS of spectral libraries downloaded from MassBank of North America using MS PepSearch (release: 02/22/2019; 0.01 Da mass tolerance for precursor and fragment searches). From the MS PepSearch output, features with the highest dot product of the same identifier were retained, and then metabolites with a dot product of <500 were removed. Furthermore, Global Natural Products Social (GNPS) Molecular Networking and Library Search were used to identify metabolites in the experimental MS/MS data ^108^. Therefore, for each feature from the metabolite data table, an MS/MS was selected based on the highest total ion count. The MS/MS with the highest count was submitted to GNPS (201 features for [+] and 361 features for [−]). The GNPS Library Search was conducted with the following settings: The precursor ion mass tolerance and fragment ion mass tolerance were set to 0.01 Da, minimum matched peaks were set to 1, and the score threshold was set to 0.5. GNPS Molecular Networking was performed with the following settings: the precursor ion mass tolerance was set to 0.01 Da, the MS/MS fragment ion tolerance was set to 0.01 Da, and the cosine score was set to >0.5 with a minimum matched peak of 1, TopK was set to 10, the maximum component size was set to 100, the maximum shift was set to 200 Da, the minimum cluster size was set to 1, and the maximum analog search mass difference was set to 500.

Bile acids were also annotated in the (−)) HILIC dataset with an error window of 0.005 Da, taking the following adducts into account: [M-H]^−^, [M+Cl]^−^, [2M-H]^−^, and [M+CH_3_CHOO]^−^. Seven bile acids conjugated with sulfate were putatively annotated as [M-H]^−^ with an error window of 0.005 Da, resulting in 53 features. To confirm the identity of bile acids, we performed an identification step with semi-targeted peak picking of bile acids, as described by Sillner et al. ^109^, with a UHPLC system (ExionLC; AB Sciex LLC, Framingham, MA, USA) coupled to a quadrupole TOF mass spectrometer (X500 QTOF MS; AB Sciex LLC). The chromatographic settings were identical to those used by Sillner et al., whereas MS detection was performed by the Q-TOF instrument in (−) ionization mode (TurboIonSpray; AB Sciex LLC). The curtain gas was set to 30 psi, ion source gases 1 and 2 were set to 45 psi, and the temperature was set to 500 °C. The MS was operated in the TOF MS and TOF MS/MS scan mode. The TOF MS analyzed molecules between 65 and 1000 Da. The ion spray voltage was set to −4500 V, CAD gas was set to 7, accumulation time was set to 0.1 seconds, the declustering potential was set to −50 V, the declustering potential spread was set to 0 V, the collision energy was set to −5 V, and the collision energy spread was set to 0 V in TOF MS mode. In TOF MS/MS mode, information-dependent acquisition (small molecule) was selected, and the following parameters were used: a maximum of 10 candidate ions, an intensity threshold of 1000 counts per second, activated dynamic background subtraction, excluding isotopes (±4 Da), and mass tolerance of ±50 mDa. The TOF MS/MS acquired data between 50 and 1000 Da, with an accumulation time of 0.025 seconds, a declustering potential of −80 V, a declustering potential spread of 0 V, a collision energy of −35 V, and a collision energy spread of 15 V; the Q1 resolution was set to unit.

Forty-five bile acids (0.05–0.4 mg/mL in methanol) and 50 µL of fecal methanolic extracts of 24 mice (two [*n* = 5], 9 [*n* = 5], 15 [*n* = 4], 24 [*n* = 5], and 30 [*n* = 5] months) were analyzed for bile acid identification. Bile acid retention times (0.05 mg/mL) were manually selected using Sciex OS Analytics (version 3.0.0.3399) with an extraction width of 0.02 Da. Raw files (.wiff2) derived from the LC–MS-based separation of bile acids were processed as described above, including chemical noise subtraction, chromatographic peak picking, chromatogram isotope clustering, and annotation of known peak function (with an rt tolerance window of 0.1 minutes and an *m*/*z* tolerance of 0.005 Da). The bile acid data (*m*/*z*, rt, and maximum intensity values for each sample) were normalized to the wet fecal weight. Pearson’s correlation was performed between the bile acid data and the (−) HILIC data subset (annotated bile acids), and features with coefficients >0.8 were considered for manual identification. Using this approach, 47 features were identified, including different adducts, and 23 features were identified as [M-H]^−^ species. Supplementary Table S4.12 summarizes all the identified and age-associated metabolites from spectral library searches and the semi-targeted identification of bile acids.

### Reconstruction of the generic metamodel

A two-step procedure was followed to obtain a metamodel for each mouse. In the first step, a generic metamodel representing the individual organs and the microbiome was assembled. In the second step, a specific metamodel of each mouse was derived by integrating the expression and metagenomics data.

In the first step (i.e., reconstruction of a generic metamodel), we joined three times the human metabolic reconstruction Recon 2.2 ^34^ representing the individual organs with a microbiome metabolic model according to their physiological interactions (Fig. 3A). Therefore, the metamodel comprised three compartments corresponding to the host tissues (brain, colon, and liver), each represented by one Recon 2.2 instance and one compartment corresponding to the microbiome. A mouse-specific metabolic reconstruction was not used, as the human reconstructions are by far the best curated, and a very high overlap in metabolic content exists between mice and humans ^110^. All compartments interfaced with each other via common exchange environments, such as the gut lumen (microbiome and colon) and the bloodstream (colon, brain, and liver). To some extent, exchanges along the bloodstream are directional, following the physiological interactions of the organs. Therefore, metabolites taken up from the diet or excreted by the colon must first pass through the liver before they can be taken up by the brain. Metabolite uptake into the brain was restricted to metabolites known to cross the blood–brain barrier (see Supplementary Table S3.7 for a list) according to published data. To compile this list of compounds, literature resources ^57,111,112^, were utilized; in addition, we selected the compounds in Recon 2.2 ^34^, along with those identified on the Virtual Metabolic Human website (www.vmh.life) whose physicochemical properties (according to the Human Metabolome Database ^113^) would allow them to cross the blood–brain barrier ^114^. For the microbiome metabolic model, all individually reconstructed MAGs were merged into a single model by combining all microbial reactions of the individual bacterial cellular compartments into a single reaction space. This merged microbiome model could then interact with the human metabolic models via the lumen exchange environment where it was located.

To better account for organ-and microbiome-specific uptake and secretion of metabolites, exchange reactions of the individual compartments were split into irreversible forward and backward directions for metabolite secretion and uptake, respectively. To model the dietary uptake of the mice, the molar concentrations of all metabolites in their diet were derived and represented in the model following an established protocol ^92^. Additionally, information on the absorption of different dietary metabolites before entry into the colon was obtained to differentiate between their ileal uptake and concentrations reaching the colon. The derived diet was then integrated into the model by adding an inflow of compounds taken up in the small intestine (i.e., the absorbed part of the diet) as a direct inflow to the bloodstream upstream of the colon. The remainder of the diet was modeled as an inflow to the colonic lumen, which would thus be available for the microbiome and the colonic compartment.

Following the merging of the human metabolic reconstruction and the gapseq-derived bacterial metabolic reconstructions and integrating of dietary uptake, several energy-generating cycles (i.e., sets of metabolic reactions encompassing human and bacterial metabolic reactions that can form ATP from ADP and phosphate without the consumption of other metabolites) were identified. The reactions involved in energy-generating cycles were screened for potential problems in reaction reversibility and adapted to prevent energy generation (see Supplementary Table S3.6 for a list of modified reactions). The final generic metamodel did not contain any energy-generating cycles. The metamodel can be found under accession MODEL2310020001 in the EBI BioModels database (https://www.ebi.ac.uk/biomodels/)^115^.

### Reconstruction of mouse/context-specific metamodels

In the second step, the transcriptomic and metagenomic data of each respective mouse were mapped to the metamodel. To achieve this, StanDep ^116^ was applied to the transcriptomic and metagenomic data to derive the core reactions for each tissue required to reconstruct context-specific models using fastcore ^35^. Transcriptomic data were preprocessed by transforming counts into fragments per kilobase of transcript per million mapped reads (FPKMs). After removing genes with at least one sample with zero detected expression or a mean FPKM below 0.1 after log_2_ transformation, FPKM values were normalized using Combat ^117^ and then transformed back to their original scale. To identify core reactions, mouse genes were mapped to their corresponding human orthologues using Ensembl Biomart ^118^; these data were then used as input for StanDep. The gene expression values of all tissues were combined into a single matrix, and tissue and age groups were used as separating categories for StanDep. StanDep was applied with “chi2dist” as the distance method and “complete” as the linkage method. After screening optimal cluster numbers, predicted core reactions remained stable when using 39 clusters (i.e., the Jaccard distance of derived core reactions for StanDep runs with increasing cluster numbers was below 0.05 using at least 39 clusters).

For metagenomic data, reads were mapped to the assembled MAGs to derive species-level counts, which were then transformed to reaction abundances by deriving a reaction contribution matrix, multiplying it by the species abundance matrix, and normalizing it to a sum of one across all reactions in a sample. A reaction contribution matrix containing reactions as rows and species in columns was obtained by setting each reaction entry to “1” if the corresponding species contained the respective reaction. Subsequently, the reaction contribution matrix was normalized to a sum of one across the reactions of each species. Similar to how core reactions were derived from gene activity, reaction abundances were used as input to StanDep with the age group of the sample as the separating factor, “chi2dist” as the distance method, and “complete” as the linkage method. Following the same procedure as for the gene activity data, 15 clusters were identified as optimal for reaction abundance data.

Because mapping the microbiome data enabled the direct identification of reaction activity, it was not necessary to map gene activity (as internally carried out by StanDep) for the microbiome data. In addition to information on gene activity and reaction abundances, metabolic exchanges between individual organs and the bloodstream previously measured in pigs were included ^119^. To this end, metabolites that had been identified to be exchanged between individual organs and the bloodstream were mapped to the corresponding metabolite identifiers in Recon 2.2. If an organ took up a metabolite, the corresponding uptake reaction was added to the core reactions, and if it was secreted, the secretion reaction was added to the core reactions. Similarly, if the kidney took up a metabolite, the corresponding outflow reaction from the blood was added to the core reactions because the kidney was not modeled explicitly. Subsequently, the core reactions for each sample and the generic metamodel were used as input for fastcore to derive a context-specific metamodel for each mouse. To run fastcore, CORPSE (https://github.com/Porthmeus/CORPSE) was used as an interface to the corresponding function of the TROPPO toolbox ^120^.

### Characterization of host-and microbiome-dependence of metabolite exchange

To characterize the metamodels of individual mice, flux ranges were identified for each reaction in the model, reactions and metabolites dependent on the microbiota were determined, and dependencies of individual host reactions on microbial reactions were identified. To identify flux ranges for each reaction in each model, FVA was performed ^121^ by maximizing and minimizing flux through each reaction using the “flux_variability_analysis()” function of CobraPy ^122^ with fraction_of_optimum = 0 (no consideration of an objective function). Because internal exchange reactions were split into irreversible forward and backward steps, they were treated separately by running FVA for each reaction while blocking the corresponding opposing direction. The FVA results were summarized by determining admissible flux ranges by subtracting minimal from maximal flux. Microbiome-dependent reactions in the host were identified by repeating the FVA but blocking each microbiome reaction. A reaction was deemed microbiome-dependent if its flux range was reduced to less than 10% of the flux range of the wild-type reaction when blocking microbiome reactions. To elucidate the metabolites exchanged between the host and the microbiota (Fig. 3B), the microbiota-dependent uptake and secretion reactions of metabolites for a given organ were counted. The number of cases of microbiome-dependent secretion was subtracted from the frequency of microbiome-dependent uptake, and a metabolite was classified as being provided by microbiota to the host or vice versa if the difference was ≥10 (20% of samples).

### Identification of reaction-level dependencies between host and the microbiota

To determine the dependencies of individual host reactions on individual microbial reactions, EFMSampler ^39^ was used to sample EFMs with each host and microbial reaction as the indicator reaction for sampling. The indicator reaction is used to define the specific reaction in a model through which EFMs should be determined. For each reaction in each metamodel, EFMSampler was run either until 10,000 EFMs were sampled or >200 seconds had elapsed. The sampling parameters were eight parallel threads for sampling EFMs and minimizing the total sum of fluxes, as the objective function when determining EFMs. For each EFMSampler run, the average flux through all reactions was recorded, along with the average frequency of occurrence of reactions in EFMs. Subsequently, the sampled frequencies were averaged across all 52 mice to obtain the average participation of reactions in EFMs containing the target reaction. This yielded a matrix in which each column corresponded to a target reaction and each row indicated the frequency at which all other reactions (including microbial reactions) occurred in EFMs containing that reaction. Thus, a non-zero value for a microbiome reaction in the column of a host reaction indicates that it occurred at least once in an EFM containing the host target reaction. Similarly, a value of “1” indicates microbiome reactions that always co-occur in EFMs of the target reaction.

To compare EFM-predicted interactions to host–microbiome correlations, scores in the interaction matrix were compared between genes and microbiome reactions with significant associations versus randomly sampled pairs of host–microbiome-associated processes. To this end, for each significant host gene-microbiome reaction association for each tissue (FDR-adjusted *p* ≤ 0.1), the reactions catalyzed by the host gene were identified, and the submatrix of the interaction matrix containing those host reactions and the microbiome reaction was determined. Subsequently, the maximal interaction score for this submatrix was determined, and these scores were collected across all significant host gene–microbiome reaction associations in a tissue. Thus, a set of “true” maximal interaction scores was derived. The same analysis was performed for randomly drawn genes associated with reactions present in the tissue and randomly selected microbiome reactions. We generated 100 such randomly drawn pairs of host–microbiome-associated processes for each tissue to obtain “random” maximal interaction scores. Then, true and randomly generated maximal interaction scores were compared using the Wilcoxon rank-sum test.

To analyze the most strongly interacting metabolic processes between the host and microbiome, an interaction between a host and microbiome reaction was assumed if the microbiome reaction occurred in at least 50% of the EFMs sampled from that host reaction across all metamodels. Then, for each host–microbiome reaction pair, we determined which metabolic subsystems they were associated with and counted each corresponding host–microbiome subsystem pair across all such pairs in the interaction matrix. The enrichment of host–microbiome subsystem pairs was then tested using Fisher’s exact test comparing for each host–microbiome subsystem pair the number of mutual interactions of reactions belonging to the host and microbiome subsystems to the frequency of interactions across the entire interaction matrix. An enrichment was assumed with an FDR-corrected *p*-value of ≤0.05, calculated using the p.adjust function in R.

### Identification of aging-regulated metabolic modules

To identify aging-regulated metabolic modules, we defined sets of reactions associated with each indicator reaction used for EFM sampling. A reaction was assumed to belong to the metabolic modules of an indicator reaction if it occurred in at least 20% of the EFMs sampled for that indicator reaction. Unlike in the analysis of reaction-level dependencies between host and microbiota, we considered both the host and microbiome components of the EFMs; thus, metabolic modules contained both host and microbiome reactions. To identify aging-regulated metabolic modules, aging-induced and aging-repressed genes for each tissue (Supplementary Tables S5.1–S5.3) were translated into the reactions with which they were associated in the metabolic model. Then, for each metabolic module, we tested whether the corresponding set of reactions was enriched for aging-induced or aging-repressed metabolic reactions using Fisher’s exact test, assuming the entire set of reactions occurring in a tissue as background. A metabolic module was considered dependent on the microbiome if it contained at least five microbial reactions. Enrichment was determined according to the subsystem annotation of each reaction in Recon 2.2 using Fisher’s exact test. For transport reactions, we also added a subsystem annotation for the transport of nucleotides (encompassing desoxyribonucleic and ribonucleic acids) and amino acids. As the underlying reaction universe for each tissue in Fisher’s exact test, all reactions occurring in that tissue were used that occurred in at least one metamodel.

### Aging brain metabolome analysis

To analyze aging-associated changes in the metabolome, data provided by Feng et al. ^51^ (Supplementary Table S2 in the referenced paper) that comprised metabolites measured across several age groups for ten anatomical brain regions were utilized. Data for three-week-old mice were excluded to avoid confounding by pre-aging trajectories. Metabolites were mapped to the IDs contained in the metamodel using the provided PubChem IDs ^123^ and information from the BiGG database ^124^. Subsequently, for each metabolite, concentrations for each measured tissue were correlated against the age of the mice using Spearman’s rank correlation. Only associations with a Benjamini–Hochberg FDR-adjusted ^125^ *p*-value of ≤0.05 were retained. In the analysis of microbiome-produced and microbiome-consumed brain metabolites, a metabolite was assumed to be provided from the microbiome to the brain if its number of cases of microbiome-dependent uptake was greater than its number of cases of microbiome-dependent secretion. An uptake or secretion reaction of the brain was assumed microbiome-dependent if its flux range was reduced to ≤10% of its wild-type value when microbiome reactions were blocked.

## Data Availability

Metagenomic raw read and MAG assembly data was deposited in the European Nucleotide Archive (ENA) under BioProject PRJEB73981 (<u>ebi.ac.uk/ena/browser/view/PRJEB73981</u>). Individual accession numbers for each MAG were listed in Supplementary Table S1.2. Gene expression data was published in the GEO database under record GSE262290 (<u>ncbi.nlm.nih.gov/geo/query/acc.cgi?acc=GSE262290</u>). Metabolomics data has been made available at the MassIVE database (<u>massive.ucsd.edu</u>) with Identifiers MSV000094409 and MSV000094410. The metamodel can be found under accession MODEL2310020001 in the BioModels database (<u>ebi.ac.uk/biomodels/MODEL2310020001</u>) ^115^. Detailed sample metadata, the microbial metabolic models and supplementary resources as well as source code used for data analysis (github.com/sciwitch/MouseMicrobiomeAging) were deposited in a zenodo record (doi.org">(doi.org/10.5281/zenodo.10844503).

## Acknowledgements

We acknowledge funding by the German Research Foundation to CK within the scope of CRC1182 (project A1.5), the Research Group miTarget (FOR5042), and the Cluster of Excellence “Precision Medicine in Chronic Inflammation” (EXC2167). We also acknowledge funding from the German Research Foundation (DFG, project number: 416 418087534) to CK and CF, the Carl Zeiss Foundation IMPULS program (project number: P2019-01-006) to CF, and the European Union’s Horizon 2020 Research and Innovation Programme (under the Marie Sklodowska-Curie grant: agreement no. 859890 [SmartAge]) to OWW, CK, and CF. This work was supported by the DFG Research Infrastructure NGS_CC (project number: 407495230) as part of the Next Generation Sequencing Competence Network (project number: 423957469). Short-read sequencing was conducted at the Competence Centre for Genomic Analysis (Kiel, Germany). We thank Katja Cloppenborg-Schmidt for the excellent technical assistance in preparing the samples for metagenomic sequencing.

## Author Contributions

Conceptualization: CF, CK, DE, LB, MH, OWW, and SS; Methodology: ASK, AW, CF, CK, DE, GM, JFB, JZ, LB, MG, MH, PSK, RH, RS, SF, SK, SW, and TD; Software: ASK, CK, GM, JT, JZ, LB, SF, SW, and TD; Data Collection: AW, CF, CK, DE, LB, MG, MH, PSK, RH, RS, and SK; Data Analysis: ASK, AW, CK, GM, JZ, LB, SF, SW, and TD; Writing: ASK, AW, CK, GM, JZ, LB, SF, and TD; Revision: CF, JFB, and JT; Visualization: CK and LB; Supervision: CF, CK, OWW, PSK, and SW; Funding Acquisition: CF, CK, JFB, and OWW; Project Administration: CF, CK, and LB. *The second authors contributed equally; their order was determined alphabetically. To accurately represent their equal contributions in any type of professional or academic documentation, these authors are permitted to rearrange the order of their names among the shared second author positions at their discretion.

## Supplement

### Study setup and microbiome characterization

**Suppl. Fig. 1:**
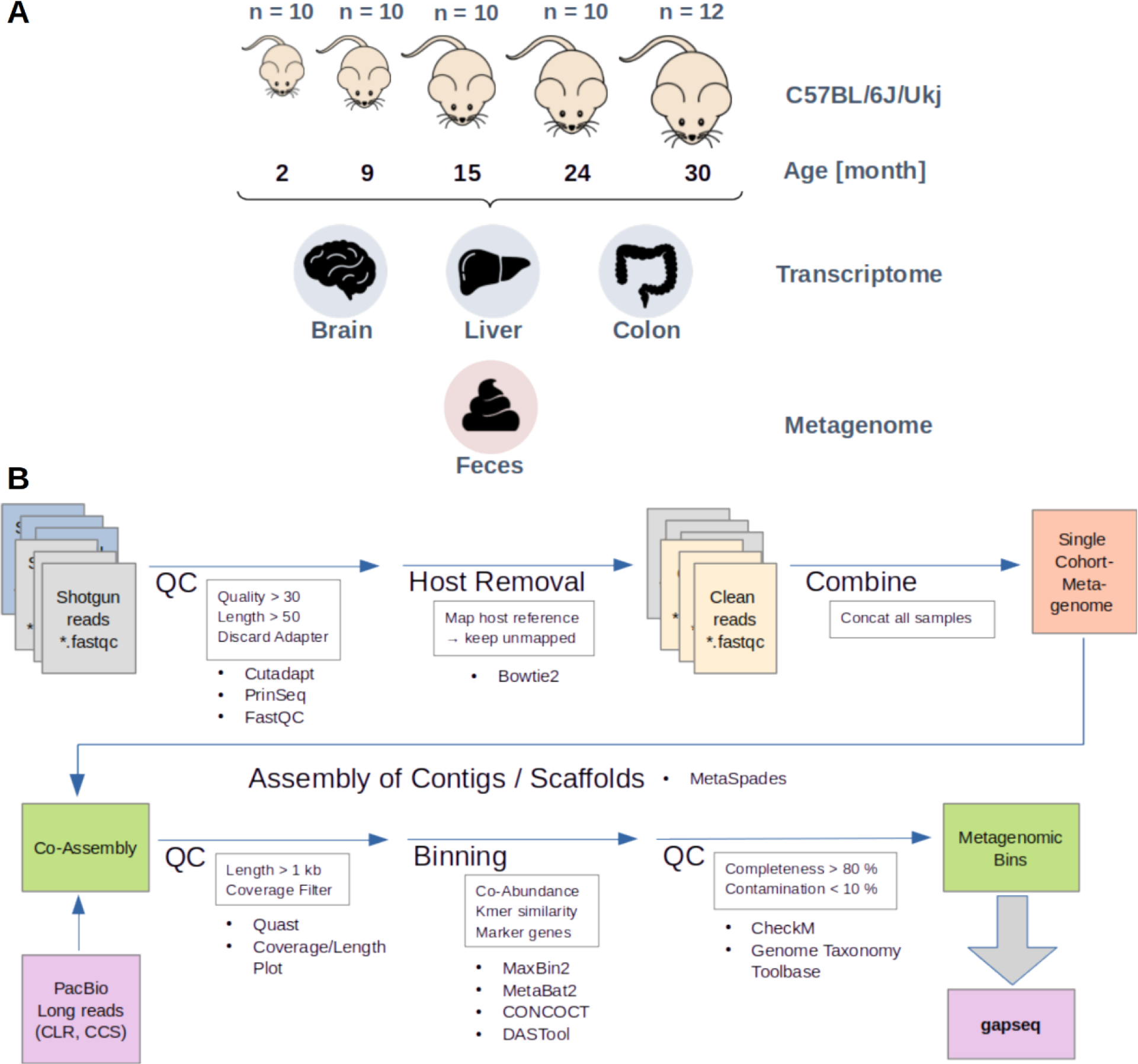
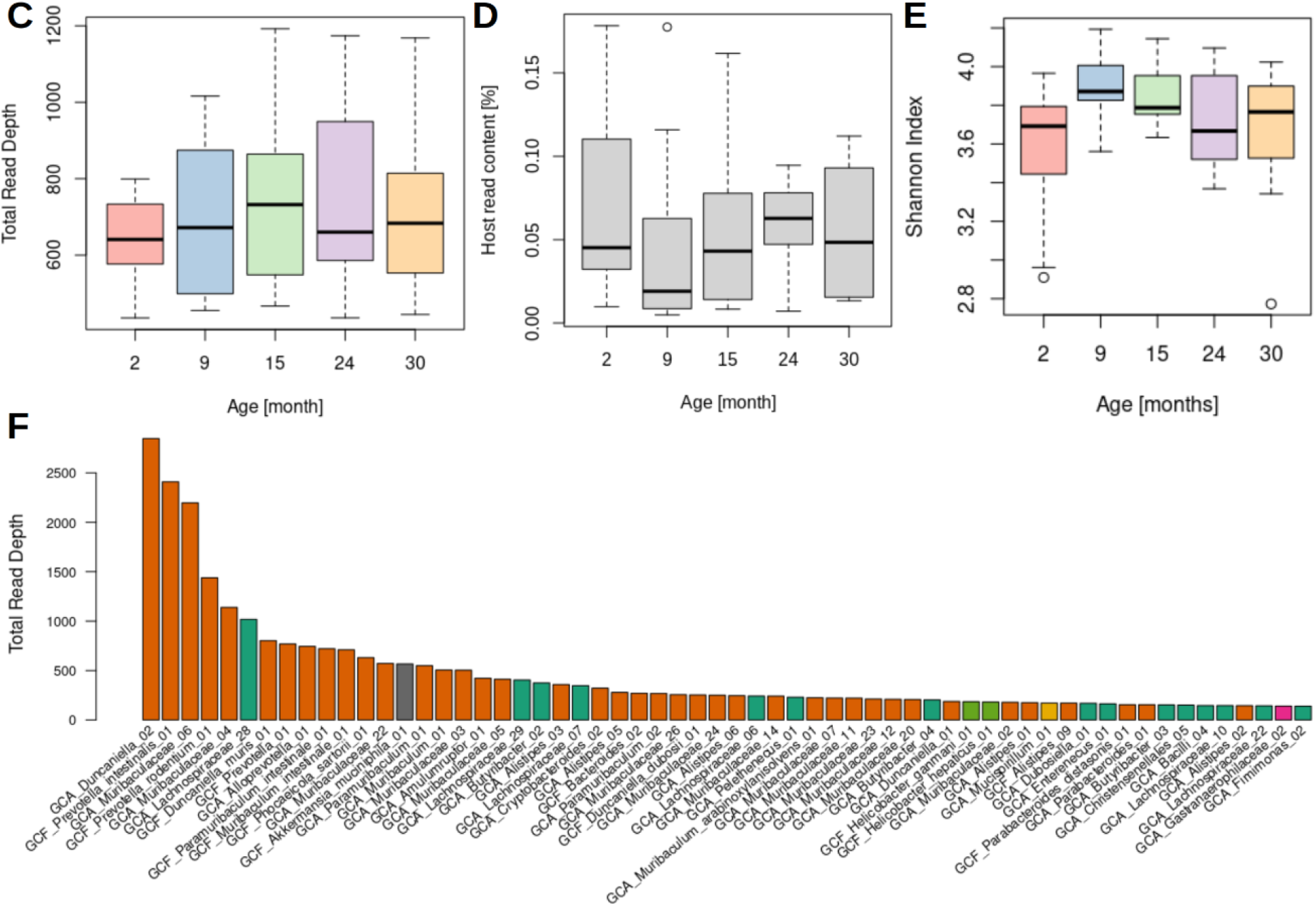
A Study setup and **B** Metagenomics workflow and Metagenome characterization. Abundance of metagenomics reads derived from microbes **C** or mouse DNA **D**. Read depths and host reads did not differ significantly between age groups according to pairwise comparisons using the Kruskal–Wallis test. **E** Alpha diversity derived from metagenomic reads mapped against MAGs. Metagenomic-derived alpha diversity did not differ significantly between the age groups according to pairwise comparisons using the Kruskal–Wallis test. **F** Abundance profile of the 60 most abundant MAGs in the full cohort. Bacteroidota predominated in the cohort while some species in Bacillota and other phyla contributed only a small amount to the total microbiome biomass. Phylum colors are reused from main Fig. 1A.

### Host–microbiome interaction

**Suppl. Fig. 2:**
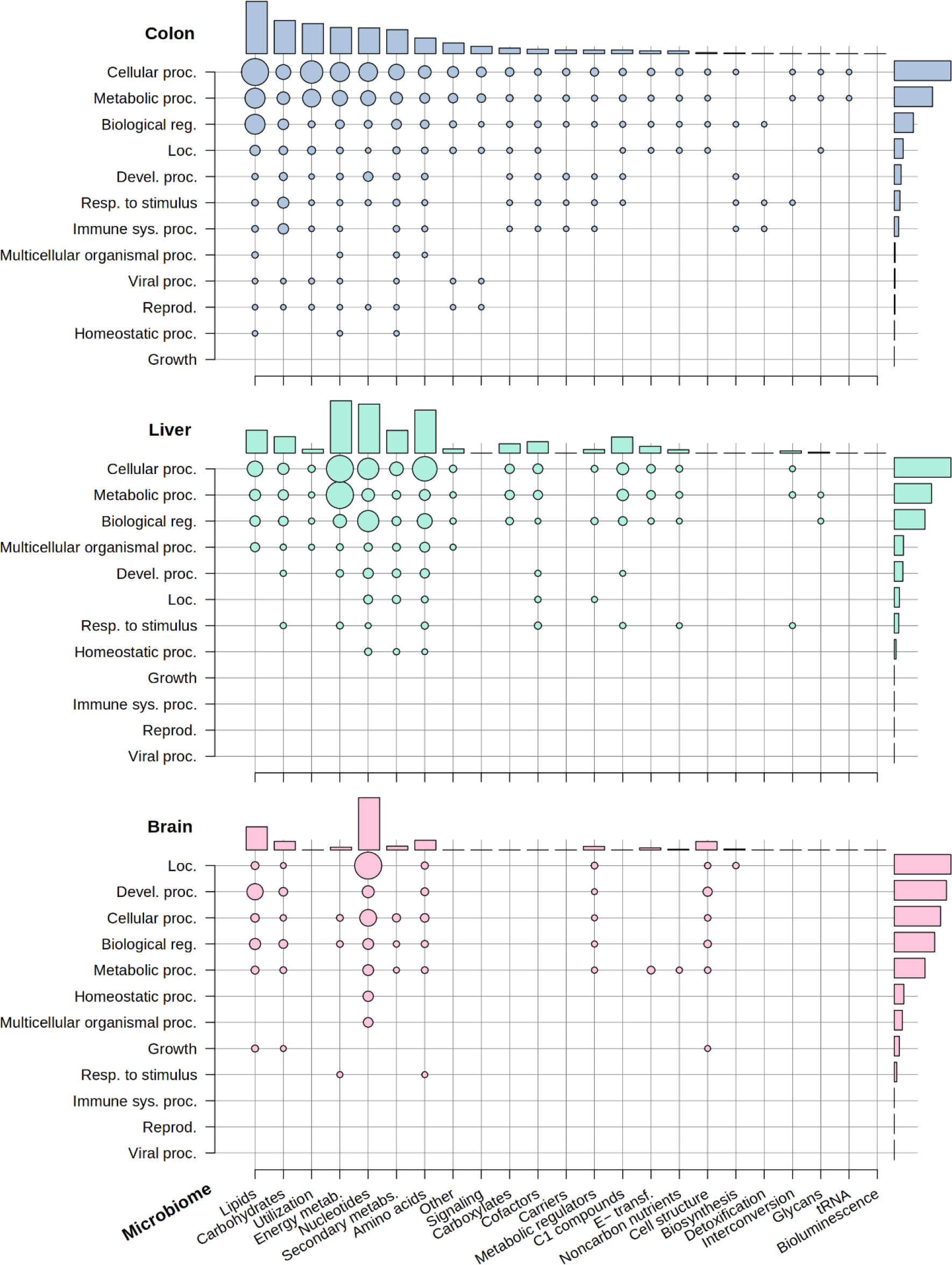
High-level pathway overview of host–microbiome abundance. Sum of −log_10_(FDR-adjusted p-values) for the odds ratios of correlation-based host–microbiome pathway interactions.

### Functional changes in the microbiome depend on host age

**Suppl. Fig. 4:**
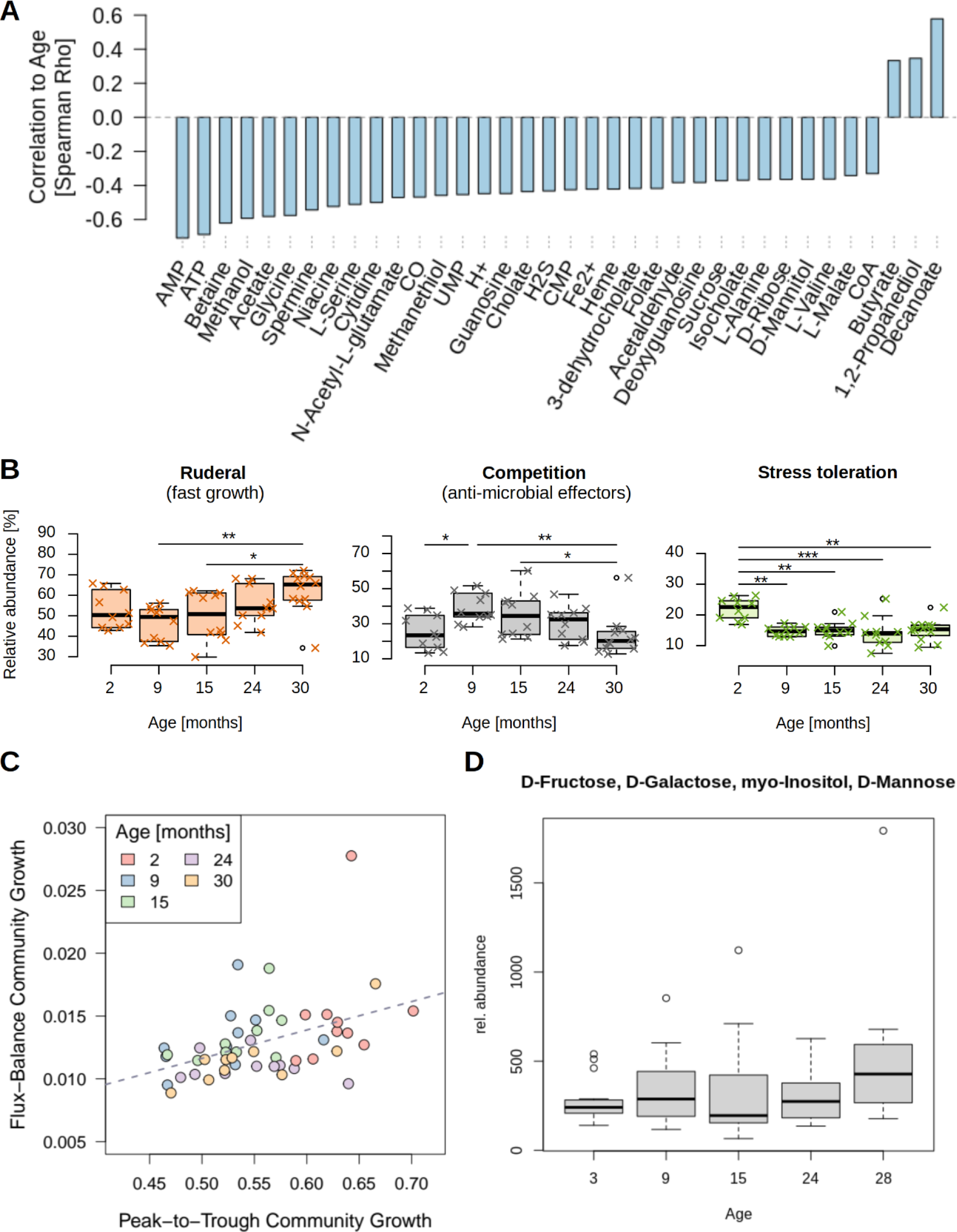
A. Age-associated changes in metabolic fluxes within the community. **B** Relative abundance of universal adaptive strategies in microbiome communities by age. We investigated the extent to which aging-associated shifts in microbiome ecology were related to species-level ecological strategies according to the universal adaptive strategies theory (UAST) framework, which categorizes species into ruderals, competitors, and stress tolerators ^44,45^. Within this framework, ruderals are species that focus on rapid growth, competitors are species that can outcompete other species through direct antimicrobial effectors and broader resource utilization, and stress tolerators are species that are resistant to stress. We observed a significant increase in the ruderal strategy with age, whereas stress tolerators and competitors significantly decreased in frequency. **C** Comparison of community growth rates derived from FBA or PTR. Growth rate estimates from metabolic modeling community simulations and metagenomic peak-to-trough analysis were strongly and positively correlated and indicated reduced growth rates of models or MAGs in older mice. **D** Changes in D-galactose concentration with age. The D-galactose concentration in mouse feces increased with age (Spearman’s ρ = 0.22, p = 0.04 [not FDR-adjusted]).

### Host aging

**Suppl. Fig. 5:**
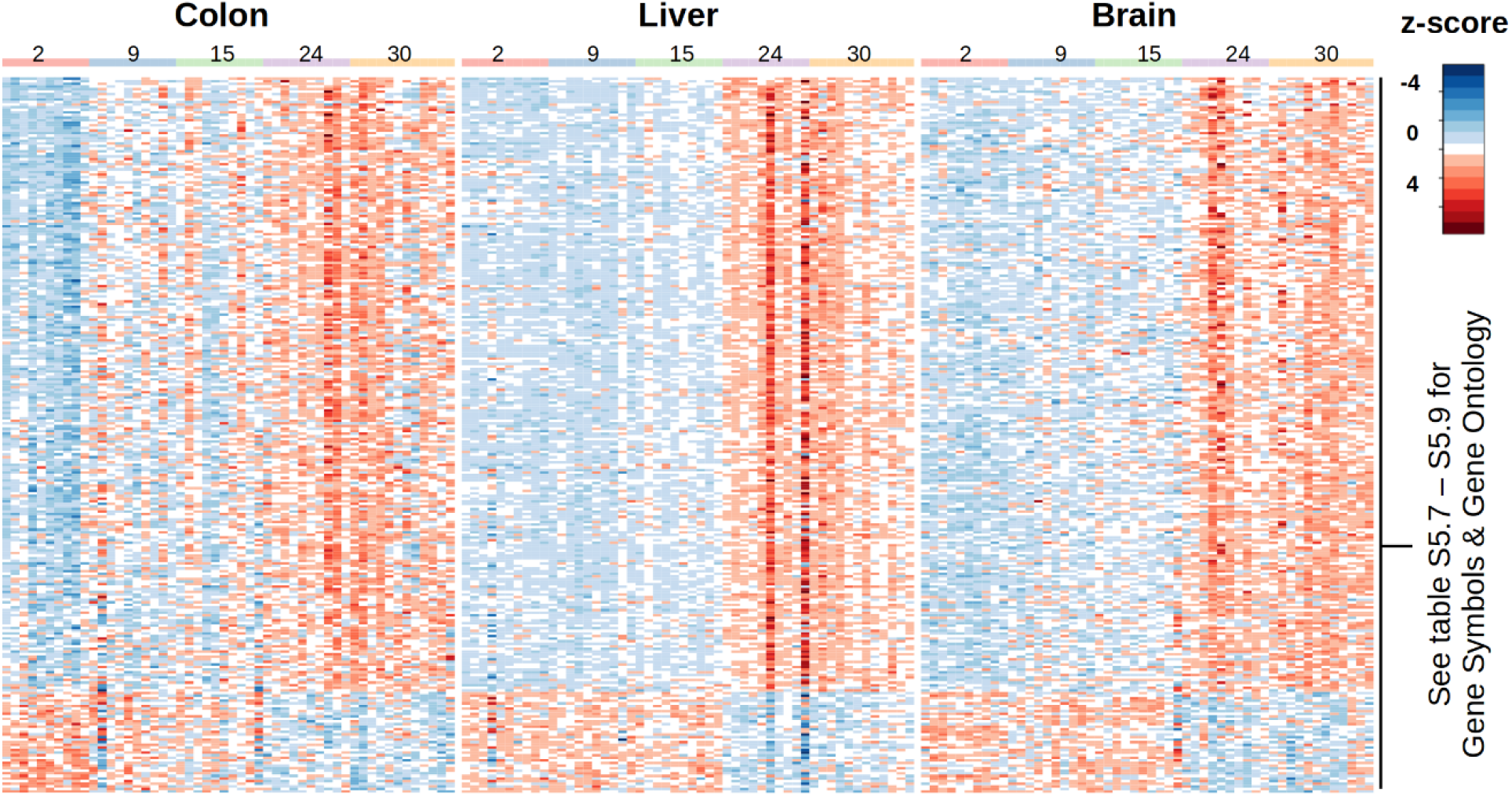
Aging-associated transcriptomic changes across host tissues. Each row represents a gene that shows common expression changes across all tissues studied (see Supplementary Table S5.7 for the complete list and Tables S5.8 and S5.9 for GO enrichment).

### Host–microbiome associations and aging

**Suppl. Fig. 6:**
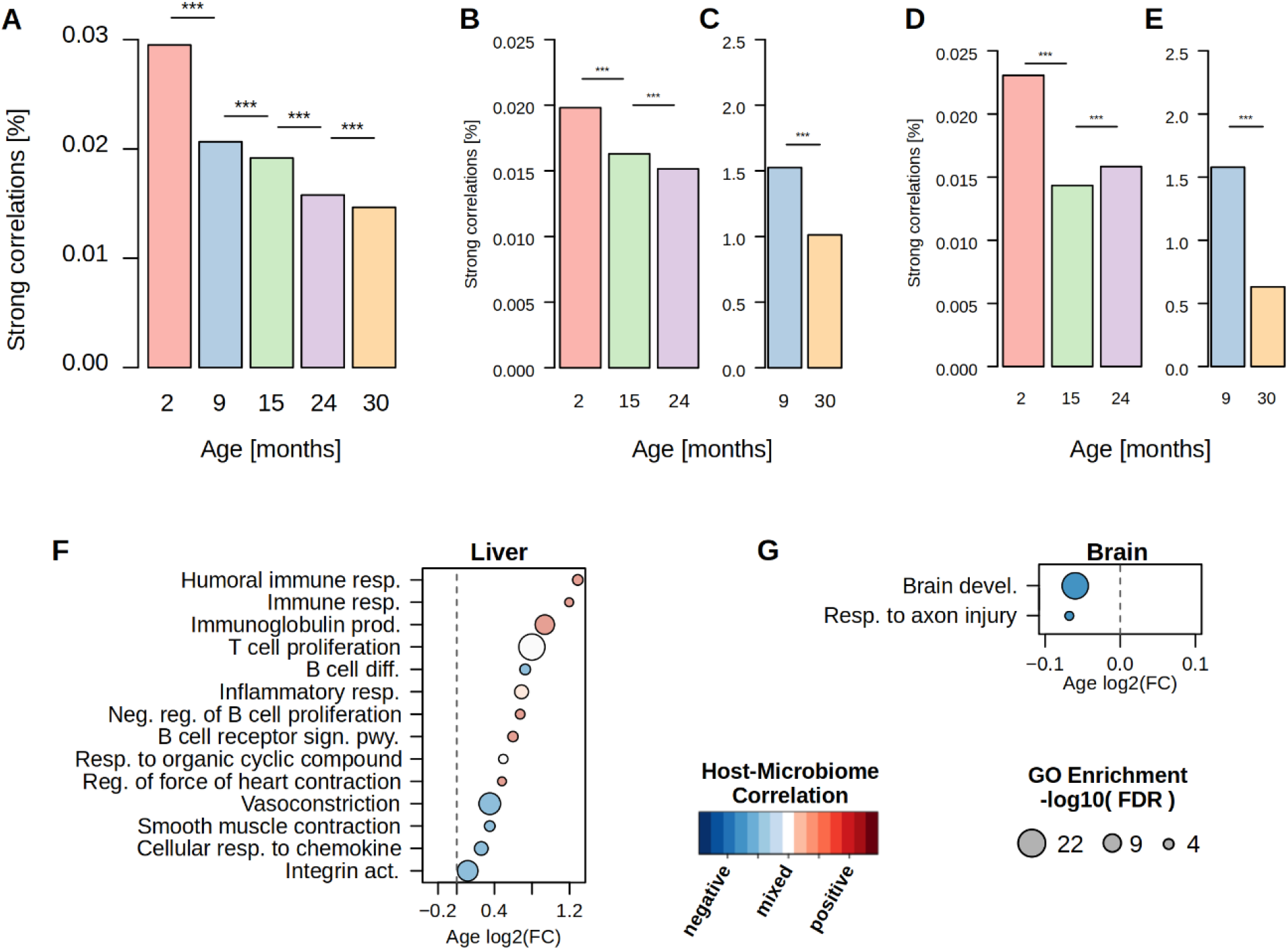
Host-Microbiome associations in aging. **A–E** Strong host–microbiome correlations stratified by age group. **A** Frequency of microbiome–colon correlations stratified by age group. Significance was tested with Pearson’s Chi-squared test with Yates’ continuity correction and Bonferroni multiple testing correction. B Liver transcripts correlated with microbiome reactions. **C** Liver transcripts partially correlated with microbiome reactions, corrected for sequencing batch. **D** Brain transcripts correlated with microbiome reactions. **E** Brain transcripts partially correlated with microbiome reactions, corrected for sequencing batch. **F–G** Organ-specific gene expression changes with age in GO biological processes that were also correlated with microbiome metabolic functions (**F** = liver, **G** = brain).

